# Golgi-Localized Mannanases Sustain Hemicellulose Biosynthesis

**DOI:** 10.1101/2025.02.25.640093

**Authors:** Talia Jacobson, Mair Edwards, Moni Qiande, Madalen Robert, Julia Moncrieff, Cătălin Voiniciuc

## Abstract

- Mannans with β-1,4-linked backbones are common cell wall components of algae and land plants. Prior challenges to enhance β-mannan content in plants point to unclear metabolic bottlenecks and the potential for hidden biosynthetic players.
- Endo-β-MANNANASEs (MANs) are the main glycosyl hydrolases that mobilize extracellular β-mannans during seed germination. However, we found that Arabidopsis *man2 man5* seeds resemble β-mannan biosynthetic mutants.
- CELLULOSE SYNTHASE-LIKE A (CSLA) overexpression restored β-mannan synthesis in the *man* double mutant and increased the distribution of crystalline polymers, but impaired the release of other mucilaginous polysaccharides.
- Using yeast synthetic biology, we dissected the functional interplay of MAN enzymes with CSLAs. Intracellular MAN2 and MAN5 reduced the quantity of insoluble β-mannan but elevated the content of water-soluble carbohydrates.
- We propose that Arabidopsis MAN2/5, and orthologous crop enzymes with a transmembrane domain, sustain hemicellulose production in the Golgi apparatus by cleaving insoluble β-mannan polymers into hydrophilic counterparts.

## Introduction

Among carbohydrate-active enzymes, glycosyl transferase (GT) and glycosyl hydrolase (GH) enzymes generally play antagonistic roles on plant cell wall polysaccharides. Hemicelluloses can cross-link cellulose microfibrils and interact with pectin domains to shape plant biology, and to define the structure of traits, dietary fibers, materials, and chemical feedstocks for industrial applications. The *Arabidopsis thaliana* (hereafter Arabidopsis) seed coat epidermal cells offer an excellent genetic model system to untangle cell wall polysaccharide biosynthesis and function. Upon contact with water, dry seeds release copious amounts of mucilaginous polysaccharides (Voiniciuc *et al*., 2015c) with smaller amounts of hemicelluloses such as branched β-mannans (Voiniciuc *et al*., 2015b) and crystalline cellulose (Griffiths *et al*., 2015). Plant β-mannan backbones are defined by their β-1,4-linked mannose (Man) residues (Voiniciuc, 2022), but can also contain β-1,4-glucose (Glc). In Arabidopsis mucilage galactoglucomannan (GGM), the Man units are substituted with single α-1,6-linked galactosyl (Gal) residues (Voiniciuc *et al*., 2015b; Yu *et al*., 2018), or β-1,2-Gal-α-1,6-Gal disaccharides (Yu *et al*., 2022). Highly branched GGM contributes to mucilage density and crystalline cellulose deposition (Voiniciuc *et al*., 2015b), while another hemicellulose, branched xylan, is important for the adherence of pectic rhamnogalacturonan-I (∼90% of Arabidopsis mucilage) to the seed surface (Voiniciuc *et al*., 2015a; Ralet *et al*., 2016). Other seed-extracted galactomannans (e.g. guar gum and locust bean gum) are frequently added as thickening agents, stabilizers, and emulsifiers in numerous food and household products, or serve as dietary fiber supplements to control appetite and reduce blood Glc levels in patients with type 2 diabetes (Juhász *et al*., 2023).

Since their backbone and branches are composed entirely of hexose sugars (Scheller & Ulvskov, 2010), plant mannans have desirable properties for a circular bioeconomy and could support the bioengineering of new products or sustainable feedstocks for biomanufacturing (Voiniciuc, 2022, 2023; Jacobson *et al*., 2024). Despite long-standing demands to elevate hexose sugar content in lignocellulosic biomass (Pauly & Keegstra, 2008), β-mannan content has been challenging to elevate in plants. For instance, seed-specific expression of a guar mannan synthase in the legume *Medicago truncatula* severely disrupted metabolism and unexpectedly decreased galactomannan levels (Naoumkina *et al*., 2008).

As recently reviewed (Voiniciuc, 2022), β-mannan biosynthesis is orchestrated by several classes of Golgi-localized glycosyl transferase (GTs) with a suite of secreted glycosyl hydrolases (GHs) expected to modify polysaccharides after their deposition in the cell wall. CELLULOSE SYNTHASE-LIKE A2 (AtCSLA2) is the chief catalyst for β-mannan elongation in the Arabidopsis seed coat (Yu *et al*., 2014). During mucilage GGM elongation, MUCILAGE-RELATED10 (AtMUCI10 also known as AtMAGT1) decorates Man residues with single α-1,6-Gal side chains (Voiniciuc *et al*., 2015b; Yu *et al*., 2018), which can be further substituted with β-1,2-Gal residues by AtMBGT1 (Yu *et al*., 2022). Plant mannans are often galactosylated (Scheller & Ulvskov, 2010) or acetylated (Zhong *et al*., 2019) because *in vitro* assays show that β-mannan oligosaccharides crystallize as they elongate (Grimaud *et al*., 2019). In the yeast *Pichia pastoris* (hereafter *Pichia*), MANNAN-SYNTHESIS RELATED1 (AtMSR1) acts as a co-factor that enables AtCSLA2 to produce glucomannan instead of relative pure mannan (Voiniciuc *et al*., 2019). Since *Amorphophallus konjac* AkCSLA3 can produce glucomannan even without MSR1 in yeast, it has been used to investigate the protein regions important for substrate specificity (Robert *et al*., 2021), to illuminate the localization and bottlenecks for hemicellulose biosynthesis (Grieß-Osowski *et al*., 2025).

Once β-mannans are deposited in the cell wall, endo-β-1,4-MANNANASES (MANs) become the chief catalysts for the backbone modification and degradation. While the biological roles of different mannan structures remains remain ambiguous (Boerjan *et al*., 2024; Guevara-Rozo *et al*., 2025), MANs are involved in wall softening and carbohydrate release during plant development (Rodríguez-Gacio *et al*., 2012). MAN hydrolytic activity increases in a wide range of plant species during seed germination, presumably to mobilize carbon reserves and to promote growth (Downie *et al*., 1997; Nonogaki *et al*., 2000; Wang *et al*., 2005; Iglesias-Fernández *et al*., 2011, 2020; De Farias *et al*., 2015; Carrillo-Barral *et al*., 2018). Four of the seven *Arabidopsis MAN* genes are expressed in dry seeds and three different single mutants (*man5*, *man6,* and *man7*, but not *man2*) decreased or delayed germination (Iglesias-Fernández *et al*., 2011). Mannan oligosaccharides released by AtMAN6 are proposed to activate signaling for xylem thickening, while AtMAN3 and AtMAN7 are induced by cadmium to increase apoplastic Man and promote heavy metal tolerance (Chen *et al*., 2015; Wu *et al*., 2023; Zhang *et al*., 2024). Some MANs also have transglycosylation activity to essentially ‘cut-and-paste’ polymers, e.g. for fruit ripening (Schröder *et al*., 2009). Intriguingly, some *MANs* are co-expressed with *CSLAs* during seed development in Arabidopsis (Voiniciuc, 2022) and in crops like coffee (Joët *et al*., 2014), but their impact on hemicellulose accumulation remains unknown (Voiniciuc, 2022). Here, we apply plant and yeast bioengineering strategies to demonstrate that Arabidopsis MAN2 and MAN5 have non-canonical roles in the synthesis of soluble β-mannans.

## Materials and Methods

The following sections describe the major materials and methods, with additional details for plant cultivation conditions, DNA construct assembly, yeast growth, biochemical analyses, cytometry and microscopy in the supplemental information (Methods **S1)**.

### Genetic Materials and Analyses

Arabidopsis T-DNA insertion mutant seeds were obtained from the ABRC or NASC stock centers, *man2-1* (SALK_126628c.20.a), *man2-2* (SALK_016450), *man5-1* (707G06.01.a), *man5-2* (SALK_015220), *man5-3* (SALK_068367), *man2-1 man5-1* (2716/1a1.01), *muci10-1* (SALK_061576), and *csla2-3* (SALK_149092). *Agrobacterium*-mediated transformations of *Arabidopsis* plants, and transient assays in *Nicotiana benthamiana* leaves were conducted using established protocols (Yang *et al*., 2022). For co-localization experiments, plasmid DNA for the Golgi yellow fluorescent protein (YFP) marker G-yk, previously validated for Arabidopsis (Nelson *et al*., 2007), was obtained from ABRC (CD3-965).

Table **S1** lists the genotyping primers for the plant Touch-and-Go Method (Berendzen *et al*., 2005) or for bacteria or yeast colony polymerase chain reaction (PCR). Cloning was mainly performed Golden Gate assembly (Engler *et al*., 2014; Marillonnet & Grützner, 2020), or NEB HiFi cloning (Table **S2**), as noted in Methods **S1**. Transcript levels in mature green siliques were analyzed using prior methods (Voiniciuc *et al*., 2015b) and normalized to the geometric mean of two housekeeping genes (Table **S1**).

### Biochemical Analysis of Mucilage and Yeast Cell Wall Fractions

Total mucilage was extracted for monosaccharide profiling via High-Performance Anion-Exchange Chromatography with Pulsed Amperometric Detection (HPAEC-PAD) as specified by (Voiniciuc *et al*., 2015b; Voiniciuc & Günl, 2016), using an established Metrohm HPAEC-PAD system and eluent gradient (Robert *et al*., 2021). *Pichia* alkaline-insoluble (AKI) polymers were isolated as previously described (Robert *et al*., 2021), followed by the separation of alkaline-soluble fractions (Voiniciuc *et al*., 2019). Water-soluble glycans were extracted from *Pichia* cells as previously described (Cocuron *et al*., 2007) with an incubation step in 70% ethanol for 5 min and a subsequent incubation in water for 5 min at 100 C without Triton X-100. After centrifugation, the supernatant was removed, hydrolyzed, and injected for HPAEC-PAD analysis. The remaining AKI was incubated in NaOH for 90 min followed by hydrolysis. Water-soluble glycans from Arabidopsis dry seeds were analyzed in a similar manner. Seeds were first ground and then incubated in ethanol and then water. The water-soluble and alcohol-insoluble residue (AIR) fractions were then hydrolyzed with trifluoroacetic acid and analyzed via HPAEC-PAD.

### Light Microscopy of Plant Materials and Yeast

Seed mucilage area was stained with ruthenium red (RR) (Voiniciuc *et al*., 2015b) and quantified using an AXIO Vert A1 or a stereoscope equipped with an Axiocam 208 color camera or a 202 monochrome camera (all from Zeiss). Seed birefringence in water was captured with an adjustable polarizer/analyzer module, and intensity plots were generated as previously described (Voiniciuc *et al*., 2016). Confocal images were captured on a Leica Stellaris 8 at the University of Florida Interdisciplinary Center for Biotechnology Research (UF/ICBR) Cytometry Core Facility (RRID: SCR_019119). Unless otherwise noted in figure legends, the 20x (0.75 NA, numerical aperture) objective was used with additional zoom in the Leica LAS X software. Calcofluor white (CF) staining of seeds (Voiniciuc *et al*., 2015b) or yeast (Robert *et al*., 2021) was performed using 0.01% Fluorescent Brightener 28 (Sigma Aldrich). Mucilage immunolabeling was followed an established protocol (Voiniciuc, 2017), using four monoclonal antibodies CCRC-M36, CCRC-M70, CCRC-M75, and CCRC-M170 (CarboSource Services, University of Georgia) and a goat anti-mouse Alexa Fluor 488 secondary antibody (ThermoFisher Scientific, A28175). Confocal images were acquired with the following (Excitation | Emission) wavelengths: CF or turquoise protein fluorescence (405 nm | 450-470 nm), green or yellow protein fluorescence (500 nm | 505-550 nm), red fluorescence (550 nm | 600-640 nm). All plant and *Pichia* images were processed and analyzed with the Fiji distribution of ImageJ (Schindelin *et al*., 2012).

### Application of MannanTrack Probe in Yeast

The MannanTrack probe used here is corresponds to + pFDH1:PaCBM35:pmScarlet-I-2A-pSmTq2(stop)ScCYC1tt αMF-PaCBM35-mScarlet-P2A-mTurquoise2 and is introduced in the Results section, and is part of an upcoming collection of genetically encoded probes for plant mannan synthases (Robert *et al*, manuscript in preparation). This MannanTrack coding sequences were codon optimized for *Pichia pastoris*, synthesized and assembled using Golden Gate cloning in the GoldenPiCS vector BB3rN_14, under the control of the pFDH1 methanol-inducible promoter and the ScCYC1tt transcriptional terminator (Prielhofer *et al*., 2017) Competent *Pichia* cells harboring the *AkCSLA3-Venus* cassette were transformed with the MannanTrack construct, integrated into the *RGI2* locus. After PCR genotyping and growth in YPM + G medium as previously described (Robert *et al*., 2021), a representative colony was selected and made competent (Lin-Cereghino *et al*., 2005). *MAN* episomal vectors were then introduced into the background. All colonies were verified via PCR genotyping before growth and further analyses specified in **Methods S1**.

### Statistical Analyses

Statistically significant changes were identified one-way ANOVA analyses performed in PAST 5 software (Hammer *et al*., 2001), or using Microsoft Excel’s t.test function assuming two-tailed distribution and equal variance of two samples.

## Results

### *man2 man5* Seeds Resemble Mannan-Deficient *csla2* and *muci10* Mutants

We focused on deciphering the roles of *MAN2* and *MAN5* (Fig. **1a**), two closely related paralogs with overlapping expression profiles (Iglesias-Fernández *et al*., 2011). We obtained an uncharacterized *man2-1 man5-1* double mutant from the GABI-DUPLO collection (Bolle *et al*., 2013), which targeted unlinked gene pairs in Arabidopsis. Real-time PCR confirmed that single and double mutants significantly reduced the transcript levels of the respective genes in developing siliques, at the stage of mucilage production (Fig. **1b**). The *man2-1 man5-1* seeds had 30% smaller mucilage capsules than the wild type (WT), (Fig. **1c**,**d**), while single *man2/5* mutants were not significantly different. Such defects are observed in several mucilage mutants, including *csla2* (Yu *et al*., 2014) and *muci10* seeds with arrested β-mannan synthesis (Voiniciuc *et al*., 2015b). In contrast to the reduced *MAN2* and *MAN5* transcript levels, the expression of *CSLA2* or *MUCI10* was not significantly altered in the *man2 man5* double mutant (Fig. **1b**). Additional insertions in *MAN5* could not be verified (Table **S1**), so we crossed an independent *man2-2* allele to *man5* plants and isolated a novel *man2-2 man5* double mutant, which also had compact mucilage compared to WT, without altering the seed dimensions (Fig. **1c-e**). Consistent with prior *man5* data (Iglesias-Fernández *et al*., 2011), only 50% of *man2 man5* seeds germinated after 48 h after stratification, compared to 80% of WT seeds (Fig. **2b**). The double mutant seeds had delayed germination across growth batches (Figs. **2b**, **S1c)**, but caught up with WT and *csla2* seeds by 96-120 h.

**Fig. 1.**
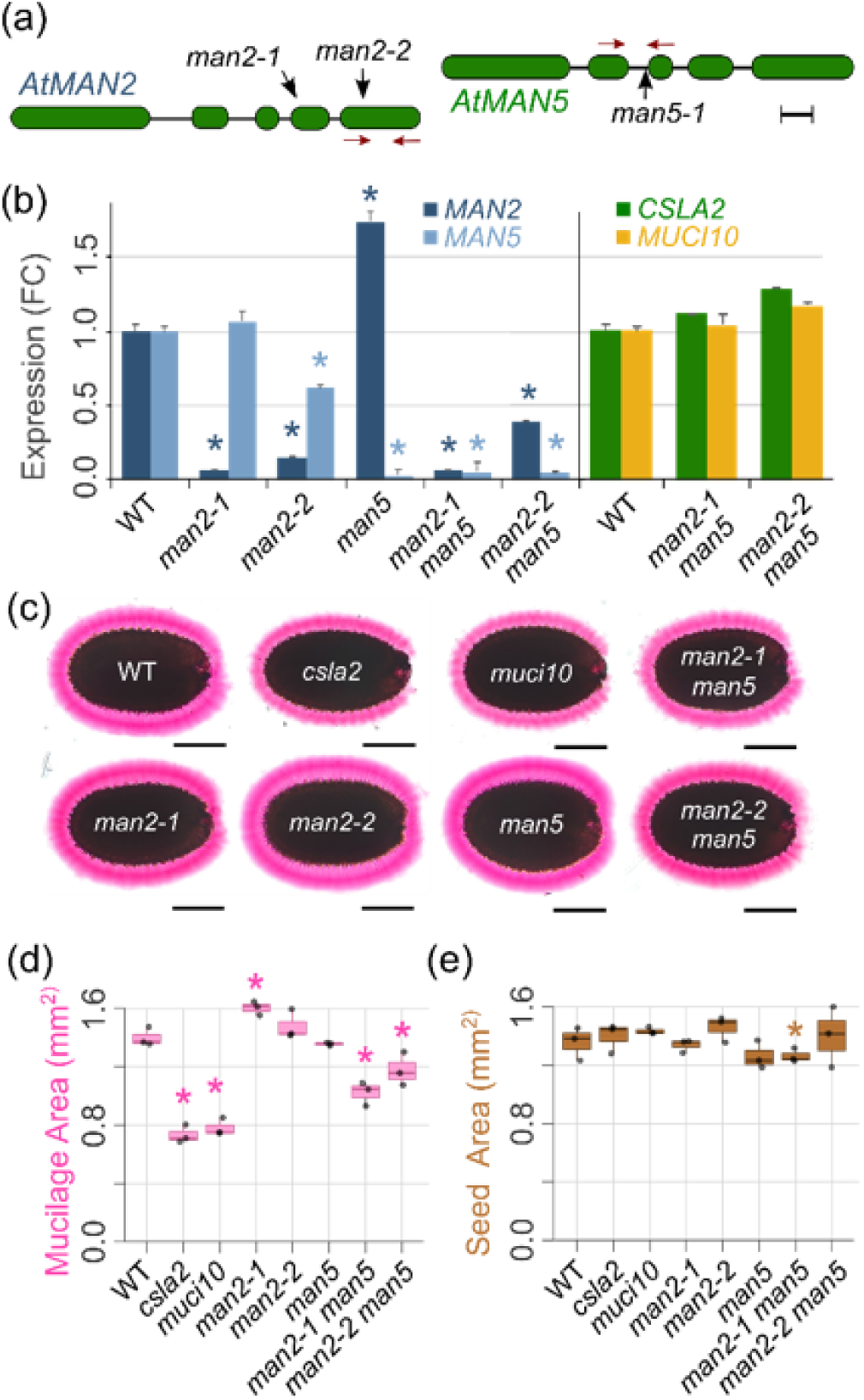
Mannanase double mutant Arabidopsis seeds have decreased mucilage accumulation. (a) T-DNA insertions in *AtMAN2* and *AtMAN5* genes, showing exon and intron structure. Red arrows indicate positions of primer binding utilized for quantitative PCR (Scale bar, 200 bp). (b) Fold changes in *AtMAN2* and *AtMAN5* or *AtCSLA2* and *AtMUCI10* transcription relative to WT in mature green siliques. Data show mean + SD of three biological replicates normalized to the geometric mean of *UBQ5* and *GAPC1*. Asterisks indicate significant decreases relative to WT (Student’s *t* test, *P* < 0.01). (c) Pectin release of WT and mutant seeds stained with RR. (d) Projected mucilage and (e) seed surface area. Data shows mean + SD of three biological replicates (around 20 seeds each). Asterisks indicate significant decreases relative to WT (Student’s *t* test, *P* < 0.05). Bars: (c) 200 μm.

**Fig. 2.**
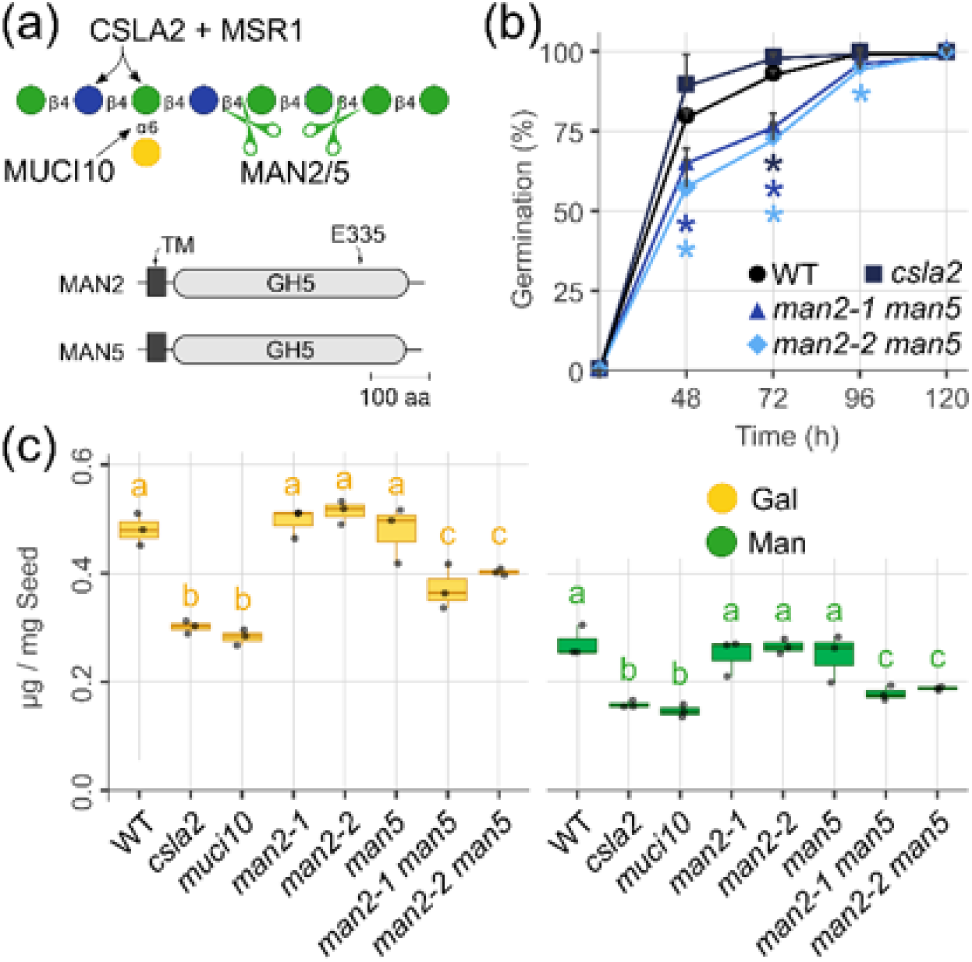
The *man2 man5* mutant seeds are impaired in β-mannan elongation. (a) Schematic of mucilage mannan biosynthesis and MAN2/5 protein domains. (b) Mean + SD germination rate for three biological replicates. For each time point, asterisks indicate significant decreases relative to WT (Student’s *t* test, *P* < 0.05). (c) Absolute monosaccharide levels of three biological replicates. Different letters denote significant changes with one-way ANOVA (Tukey’s post-hoc test, *P* < 0.01).

To understand why mutations in β-mannan synthases and hydrolases resulted in similar staining defects, despite their antagonistic biochemical activities, we profiled total mucilage composition with HPAEC-PAD. Compared to *man* single mutants and WT, *man2 man5* mucilage contained significantly less Gal and Man, the most characteristic monosaccharides of branched β-mannans (Fig. **2c**; Table **S3a**). Glc, the third component of GGM, showed similar trends but had a high coefficient of variation because it can be derived from several cell wall polymers as well as starch. Although pectin immunolabeling with CCRC-M36 showed results consistent with the RR staining defects, β-mannan antibodies (M70, M75, or M170) in the CCRC collection (Thorne *et al*., 2023) could not bind WT or *man2 man5* mucilage (Fig. **S1e**). Nevertheless, *man2 man5* mucilage phenotypes and chemotypes approached the severity of *csla2* and *muci10*, with up to 50% reductions in Gal and Man compared to WT across multiple growth batches (Figs. **2c**, **3c, S1**). These data are consistent with the participation of *MAN2* and *MAN5* in β-mannan production, rather than its catabolism in the seed coat.

**Fig. 3.**
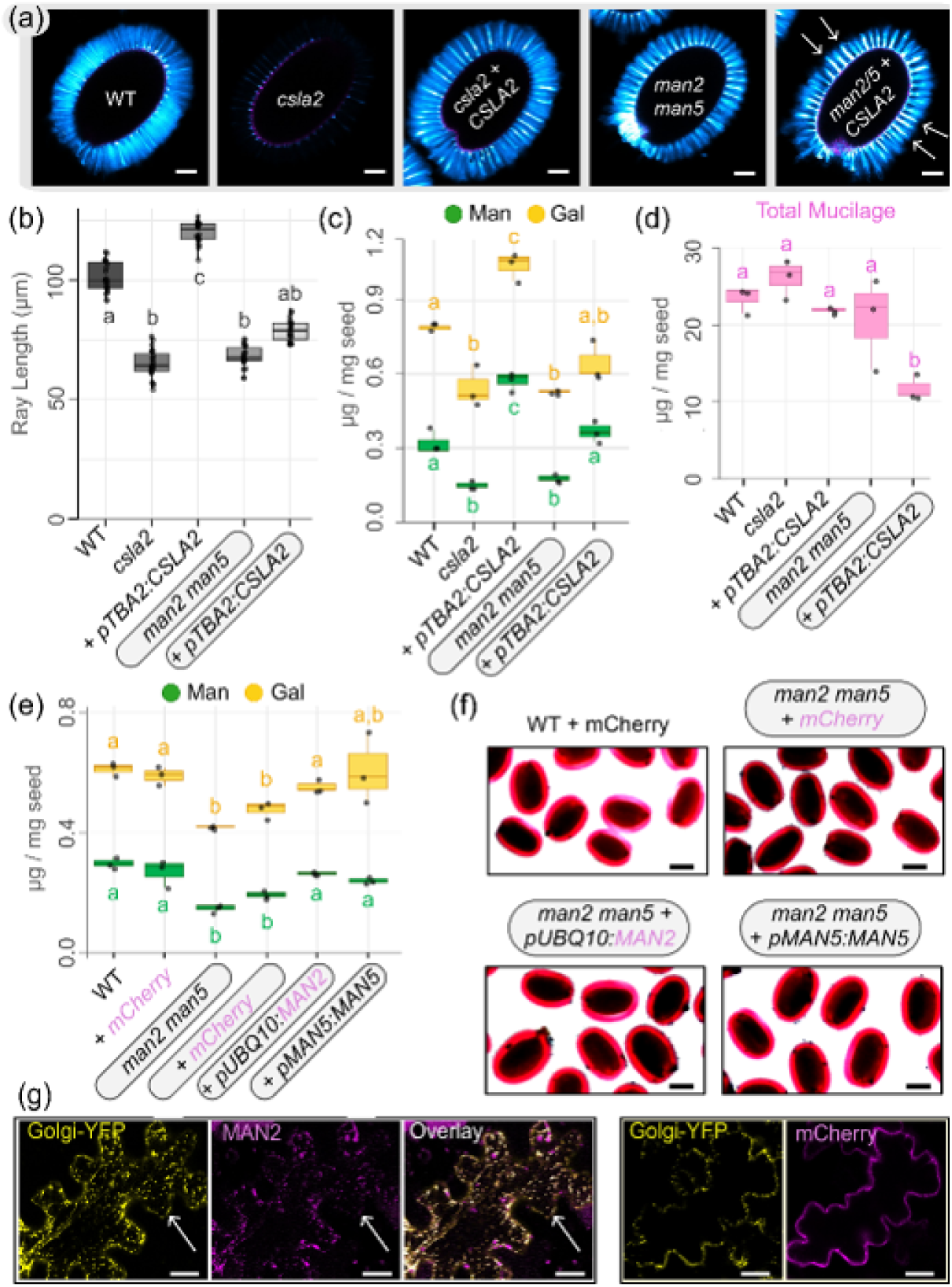
Golgi-localized MAN enzymes are important for mucilage structure. (a) CF staining of β-glucans in seed mucilage. Arrows mark disordered rays. (b) Length of CF-stained rays. (c) *AtCSLA2* overexpression affects Gal and Man content, and (d) total mucilage release. Biochemical (e) and mucilage RR staining (f) complementation of *man2 man5* by AtMAN gDNA constructs. Different letters denote significant changes with one-way ANOVA (Tukey’s post-hoc test, *P* < 0.01). (g) Optical sections of *N. benthamiana* leaves expressing a Golgi-YFP marker and AtMAN2-mCherry or an mCherry control. Arrows mark co-localization. Bars: (a) 100 788 µm, (f) 200 µm, (g) 30 µm.

### *AtCSLA2* Overexpression Rescues *csla2* Defects, but Impairs *man2 man5* Mucilage

Despite the yeast activities described later in the Results section, CSLAs and MANs cDNA sequences (Fig. **S1a,b**) have been difficult to express in plants. In this study, we found that *csla2* mucilage β-mannan deficiency (Table **S3b**) can be rescued with a seed coat-specific *pTBA2:CSLA2* construct, featuring the *TESTA ABUNDANT2* promoter (McGee *et al*., 2021). The *CSLA2* transgene restored the RR staining of pectin (Fig. **S2a)** and CF staining of cellulosic rays (Fig. **3a**) around *csla2* seeds. In contrast, when introduced in the *man2 man5* background, *pTBA2:CSLA2* led to longer yet disordered rays (Fig. **3a,b**) and a patchy mucilage capsule (Fig. **S2a**). The trends in cellulose staining were supported by imaging of crystalline polymer birefringence under polarized light. The *man2 man5* double mutants had an intermediate reduction of birefringence compared to *csla2* seeds (Fig. **S2b-d**). Although *CSLA2* overexpression in the *man2 man5* seed coat elevated Man and Gal content to nearly WT levels (**Fig. 3c**), it decreased the amount of total extractable mucilage by nearly 60% (Fig. **3d**, Table **S3b**). This was likely due to reduced mucilage release/extraction, rather than impaired secretion, because scanning electron microscopy (SEM) did not reveal major changes in the surface morphology of the mucilage pockets or the columellae (Fig. **S1d**).

### Non-Canonical MAN2 and MAN5 Localize in the Golgi Apparatus

Unlike most plant and microbial MANs, AtMAN2 and AtMAN5 are predicted to have an N-terminal transmembrane domain instead of a signal peptide (Figs. **2a****, S3**). However, determining their subcellular localization *in planta* was not straightforward and required extensive troubleshooting (summarized in Table **S4**). *AtMAN2/5* cDNA sequences did not show any detectable protein expression/function *in planta* using *pTBA2*, the putative *pMAN2* promoter, or the viral *p35S* promoter (Fig. **S1a-d**). Similar vectors worked for fluorescent protein controls and cDNA sequences for other enzymes (e.g. CSLA2), but plant gene expression may require distal or intron-localized enhancers (Vollen *et al*., 2024), which are difficult to predict. To circumvent these unexpected challenges, we amplified *AtMAN^gDNA^* genomic sequences for C-terminal *mCherry* fusions driven by the Arabidopsis *UBIQUITIN10* promoter (*pUBQ10)*. At long last, the *MAN2^gDNA^*-*mCherry* combination showed detectable fluorescence after transient expression in *N. benthamiana*. Unlike the cytosolic mCherry control, MAN2-mCherry co-localized with the known Golgi-YFP marker G-yk (Nelson *et al*., 2007), in small punctae (Fig. **3g**) and showed similar trajectories (Video **S1**), consistent with roles in mannan biosynthesis. Since *MAN5^gDNA^-mCherry* did not show any detectable fluorescence, we cloned an even longer genomic *MAN5* construct that features its putative promoter, native stop codon, and its 3’ untranslated region. Eight *pMAN5:MAN5^gDNA^* and four *pUBQ10:MAN2^gDNA^-mCherry* transformants rescued the *man2 man5* biochemical defects and compact mucilage phenotype (Fig. **3e,f**, Table **S3c**).

### Intracellular MAN2 and MAN5 Reduce Insoluble β-Mannan Accumulation in Yeast

*MAN^cDNA^-mCherry* constructs were expressed in a modular *Pichia* platform via an episomal vector compatible with Plant Modular Cloning (MoClo) standards (Grieß-Osowski *et al*., 2025). AtMAN2 and AtMAN5 proteins were localized in intracellular compartments (Figs. **4b**, **S4d**) and did not alter the glycan composition of WT X-33 cells (Fig. **4a**). While episomal expression of AtMSR1 enhanced glucomannan production in CSLA yeast strains (Fig. **4a**, **S6a**), co-expression of AtMAN2/5 with either AkCSLA3 or AtCSLA2 reduced insoluble β-mannan content by 30–40% (Figs. **4a**, **S4a**). To ensure that GH activity led to these differences, AtMAN2’s catalytic nucleophile (Wang *et al*., 2015) was mutated to an alanine (Figs. **2a**, **S3a-c**). AtMAN2^E335A^ had similar fluorescence intensity and subcellular localization to its WT sequence (Fig. **S3d**) but no longer altered mannan content (Fig. **4a**, **S4a**). Similar biochemical changes to the episomal constructs were observed for integrative *AtMAN-mRuby2* constructs driven by a weaker promoter. After alkaline treatment, the integrative *AtMAN2/5* constructs reduced the content of insoluble mannans (Fig. **S5**), without significant changes in the alkaline-soluble fractions that are rich in mannosylated yeast proteins.

**Fig. 4.**
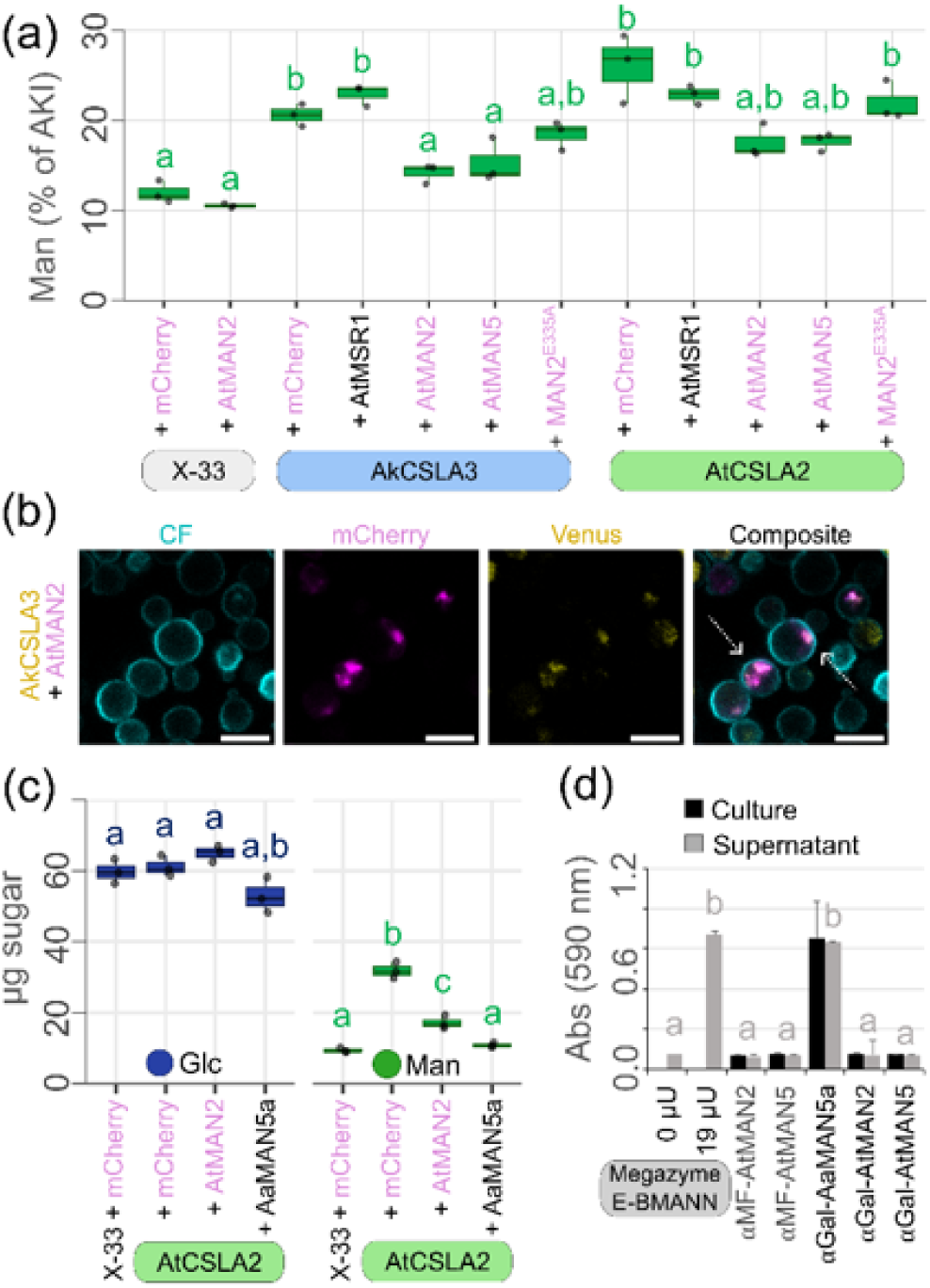
MAN2/5 enzymes reduce insoluble mannan content in yeast. (a) Relative Man content of insoluble yeast polymers. Box plots show three biological replicates. (b) Arrows mark the co-localization of AkCSLA3-Venus and AtMAN2/5-mCherry and counterstained with CF in confocal micrographs acquired with an 86x (1.2 NA) objective. (c) AKI polymers made by AtCSLA2 plus AtMAN2 or the fungal AaMAN5a. (d) AZCL-galactomannan digestion assays using *Saccharomyces* cell mixtures and supernatant, alongside commercial MAN controls (Megazyme, E-BMANN). Different letters denote significant changes with one-way ANOVA (Tukey’s post-hoc test, *P* < 0.01). Magenta labels indicate a C-terminal protein fusion to mCherry. Bars: (b) 5 µm.

### AtMAN2 Co-Localizes with AkCSLA3 and Cannot Digest Extracellular β-Mannans

Since AkCSLA3 tolerates C-terminal fluorescent tags (Grieß-Osowski *et al*., 2025), we co-expressed AtMAN2-mCherry with AkCSLA3-Venus in *Pichia* cells. Although only a subset of AkCSLA3-Venus cells showed mCherry fluorescence due to the greater heterogeneity of episomal plasmid expression, the two proteins co-localized in the endomembrane system (Fig. **4b**, **S4c,d**). Furthermore, AkCSLA3-Venus fluorescence remained constant with either mCherry or MAN-mCherry protein (Fig. **S4c**). We hypothesized that unbranched glucomannan production by CSLAs would perturb yeast cell ultrastructure, and that AtMAN2/5 enzymes could alleviate such defects. However, transmission electron micrographs of 3-day-old *Pichia* cells showed enlarged vacuoles and signs of autophagy, even without CSLAs (Fig. **S7a,b**). Although the same harvest timepoint was suitable for live cell imaging and biochemical analyses in this study and prior work (Grieß-Osowski *et al*., 2025), we could not observe Golgi bodies in thin sections (100 nm) of cells preserved via high-pressure freezing or chemical fixation. However, we noticed that AkCSLA3 cells (accumulating glucomannan) had 30% thicker cell walls compared to the negative control (**Fig. S7c,d**). Co-expression of AkCSLA3 with the wild-type AtMAN2 (but not the AtMAN2^E335A^ variant) restored the wild-type thickness.

Plant MANs are reported to have low catalytic activity (Wang *et al*., 2015) relative to microbial MANs (Ishii *et al*., 2016), such as *AaMAN5a* from the plant pathogen *Aspergillus aculeatus*. While AtMAN2 reduced insoluble Man levels in the *Pichia* AtCSLA2 strain by 46%, fungal AaMAN5a led to a 66% decrease (Fig. **4c**), resulting in WT-like yeast cell walls. Since AaMAN5a has been used to ferment β-mannans in *Saccharomyces cerevisiae* (Ishii *et al*., 2016), we also introduced *AtMANs* in this host and tested their ability to digest exogenous Azurine Cross-Linked (AZCL)-galactomannan. Blue-dyed β-mannan oligosaccharides were released by the supernatant fraction of cells secreting AaMAN5a (Fig. **4d**), or by minute amounts of a fungal MAN purified from *Aspergillus niger*. Neither AtMAN2 nor AtMAN5 could digest extracellular galactomannan (Fig. **4d**), despite replacing their N-terminal transmembrane regions (Fig. **S3a**) with the yeast α-galactosidase (αGal; enabling AaMAN5a secretion) or the α-mating factor (αMF) signal peptide. This is likely due to a lack of AtMAN protein secretion, because fluorescently tagged AtMAN2/5 proteins were retained inside *Pichia* cells despite the addition of the αMF signal peptide (Fig. **S3d**) or if their transmembrane domains were simply removed (Fig. **S4b**). However, when removing the first 34 amino acids without adding a signal peptide, MAN2(34)-mRuby2 proteins became cytosolic and no longer altered the production of insoluble mannans (Fig. **S6a,b**), which occurs in the Golgi lumen.

### Tracking Mannan Accumulation and Hemicellulose Solubility

To further quantify how AtMANs influence hemicellulose accumulation in yeast, we applied a novel, genetically encoded probe featuring a carbohydrate-binding module (CBM) derived from a microbial MAN to specifically track β-mannans. The potential utility of CBM-based probes for live cell imaging was previously reviewed (Voiniciuc *et al*., 2018b), and the development of a collection of MannanTrack probes to monitor the activity of mannan synthases in yeast and plants will be fully characterized in a separate manuscript (Robert *et al*, in preparation). Here, we briefly introduce and demonstrate the utility of one MannanTrack probe that consists of the *Podospora anserina* CBM35, which is involved in mannan-binding based on structural and biochemical analyses (Couturier *et al*., 2013). An N-terminal αMF signal peptide targets the PaCBM35 for secretion, and a C-terminal tags containing mScarlet-I (Bindels *et al*., 2017) and a superfolder mTurquoise2 ox (Meiresonne *et al*., 2019) as ratiometric reporters for red and blue fluorescence (Fig. **5a**). The two fluorescent proteins are separated by a “self-cleaving” P2A linker from the porcine teschovirus-1 2A, which can result in ribosome skipping during translation to generate separate polypeptides from bi-cistronic vectors (Liu *et al*., 2017). We observed successful skipping and recommencement of translation for our MannanTrack probe resulting in two separatee fluorescent reporters: the mannan-binding αMF-PaCBM35-mScarlet was fully secreted from *Pichia* cells without AkCSLA3 (**Fig. 5a**, additional images in **Fig. S8**), while the mTurquoise2 (downstream of P2A) remained in the cytosol.

**Fig. 5.**
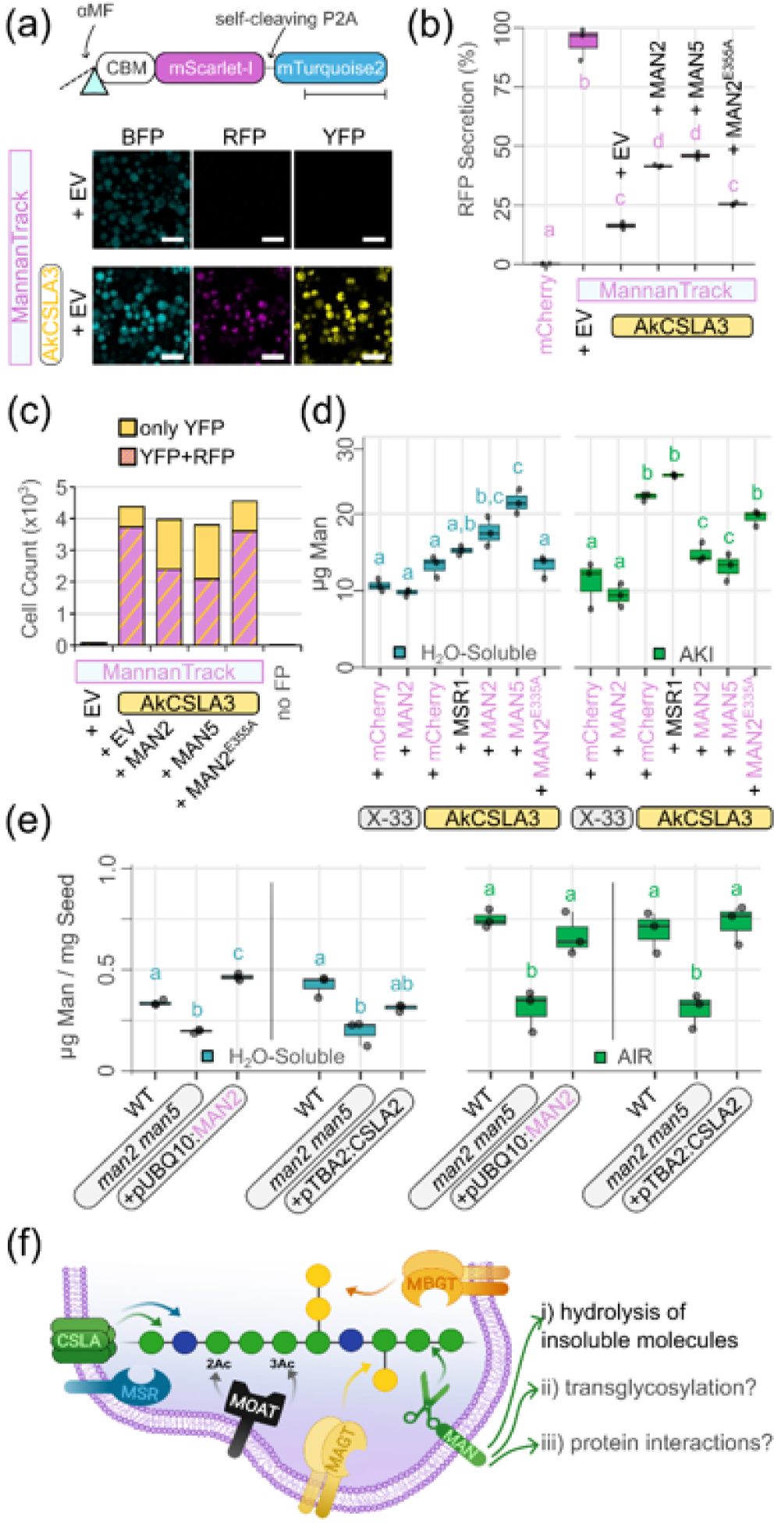
*MAN2* and *MAN5* promote mannan solubility and secretion. (a) MannanTrack probe components in *Pichia* control strains to validate the blue (BFP), yellow (YFP) and red (RFP) protein fluorescence patterns. (b) Relative secretion (supernatant RFP / cell mixture RFP) of the PaCBM35-mScarlet, normalized to the BFP cell fluorescence relative to empty vector (EV). (c) Number of cells with detectable AkCSLA3-Venus and PaCBM35a-mScarlet fluorescence quantified via flow cytometry. (d) Man content of hot water (H_2_O-soluble) and AKI polymers in *Pichia* extracts. (e) Man content of hot water (H_2_O-soluble) and AIR polymers in Arabidopsis ground seeds. Box plots show three biological replicates and different letters denote significant changes with one-way ANOVA (Tukey’s post-hoc test, *P* < 0.01). Magenta labels indicate a C-terminal protein fusion to mCherry. (f) Updated model of β-mannan biosynthesis in the Golgi apparatus (based on Voiniciuc, 2022) with AtMAN2/5 role(s). Bars: (a) 5 μm.

In contrast to yeast cells with native polysaccharide composition, the αMF-PaCBM35-mScarlet probe was largely retained inside cells (ie. reducing the RFP secretion) producing insoluble mannan (Fig. **5a,b**) and co-localized with AkCSLA3-Venus (Figs. **5a**, **S8**). Based on plate reader measurements (Fig. **5b**), confocal microscopy (Fig. **S8**), and flow cytometry (Fig. **5c**), the introduction of *MAN2/5* significantly increased MannanTrack secretion. For instance, *MAN2/5* enhanced mScarlet fluorescence in the culture supernatant by 20% relative to empty vector or *MAN2^E335A^* controls (Fig. **5b**), which prompted us to evaluate β-mannan solubility. Using hot water to directly extract soluble β-1,4-linked glucans from ground yeast cells (Cocuron *et al*., 2007), we isolated 30–60% more Man from cells co-expressing *AkCSLA3* with *AtMAN2*/5, compared to *AkCSLA3* alone or the non-functional *MAN2^E335A^* (Fig. **5d****, S9**). To determine whether MANs function similarly in Arabidopsis, we ground dry seeds and isolated alcohol-insoluble residue (AIR) as well as water-soluble fractions for HPAEC-PAD analysis (see extraction scheme in **Fig. S9**).

The *man2 man5* mutants contained significantly less Man in both fractions compared to WT, and *CSLA2* overexpression increased Man content primarily in the insoluble polymer fraction (Fig. **5e**). Therefore, we propose that MAN enzyme activity in the endomembrane system converts insoluble mannan polysaccharides into shorter and more hydrophilic molecules.

## Discussion

### Breaking A Few Bricks to Build a Wall?

Thus far, plant MANs are reported to degrade or cut-and-paste extracellular β-mannans (Bewley, 1997; Moreira & Filho, 2008; Schröder *et al*., 2009), and typically have antagonistic consequences to Golgi-localized (gluco)mannan synthases (Voiniciuc, 2023). In this study, we discovered that Arabidopsis seeds have reduced mannan production without *MAN2* and *MAN5* (Fig. **1d**, **2c, 5e**), despite wild-type *CSLA2* and *MUCI10* transcript levels (**Fig. 1b**). The significant loss of mannan accumulation in seeds (**Fig. 5e**) despite the normal secretion of other mucilage polysaccharides (**Fig. S1d,e**) implies that *man2 man5* double mutants primarily disrupt mannan elongation. Therefore, we hypothesize that intracellular MAN2/5 enzymes cooperate with Golgi-localized GTs to sustain mannan production (**Fig. 5f**). Although this represents a novel role for endo-MANs, an endo-glucanase is known to participate in cellulose biosynthesis. KORRIGAN1 (KOR1) is a plasma membrane-localized cellulase that associates with the Cellulose Synthase Complex (Vain *et al*., 2014). While genetic studies demonstrate its importance for cellulose synthesis, KOR1’s precise mode of action remains elusive. For instance, KOR1 is proposed to relieve mechanical stress placed on the cell as crystalline polymers are deposited into the wall (Mielke *et al*., 2021). Since unbranched β-mannans also form crystalline structures as they elongate (Grimaud *et al*., 2019), MAN2/5 could be needed to reduce intracellular polymer aggregates and mechanical strain (Fig. **5f**). This raises the question: why would plant cells build polysaccharides only to immediately degrade them? One possible explanation is that CSL polysaccharide synthases may lack an intrinsic mechanism to stop polymer elongation, making controlled cleavage by GHs essential for regulating fiber length and preventing aggregation.

Although *AtCSLA2* overexpression using *pTBA2* forced the elongation of mannans, it could not fully restore the content of water-soluble mannans in seeds (**Fig. 5e**) and disrupted the organization of other mucilage polymers (**Fig. 3a,b**, **S2a,d**). In the *csla2* background, *pTBA2:CSLA2* nearly doubled the amount of mannan compared to WT plants (**Fig. 3c)**. In contrast, *pTBA2:CSLA2* increased the mannan content of *man2 man5* mucilage extracts only to WT levels but significantly disrupted the release of pectin (**Fig. 3d, S2**). The *man2 man5 + CSLA2* seeds could have reduced secretion of rhamnogalacturonan-I to the cell wall and/or impaired mucilage release due to altered matrix composition and elevated content of birefringent polymers. The second option appears more likely as the SEM of *man2 man5* seeds resembled the WT (Fig. **S1d**), instead of the severe defects of seeds deficient in pectin synthesis (Voiniciuc *et al*., 2018a). Mutants such as *mum2* are known to release minimal mucilage due to elevated polysaccharide cross-linking rather than a reduction in pectin content (Dean *et al*., 2007; Voiniciuc *et al*., 2015c). Therefore, our results reinforce the importance of mannans for matrix polysaccharide architecture as well as cellulose deposition (Voiniciuc *et al*., 2015b; Yu *et al*., 2018; Voiniciuc, 2022). MAN2/5 also have functions beyond the seed coat, with similar biosynthetic roles proposed by an independent study of *man2 man5* xylem vessels (Kikuchi *et al*., 2025). We also found that *man2 man5* double mutants had delayed germination (Fig. **2c**), consistent with *MAN* gene expression in the endosperm (Iglesias-Fernández *et al*., 2011).

### How Hemicellulose Structure Impacts Matrix Polysaccharide Secretion

Despite increasing similarities between branched β-mannan and xyloglucan biosynthesis (Yu *et al*., 2022; Grieß-Osowski & Voiniciuc, 2023), major questions remain about the precise functions of different hemicellulose structures in plants (Boerjan *et al*., 2024). Xyloglucan mutants lacking Gal decorations have severe growth penalties and secretion defects (Hoffmann & McFarlane, 2024), yet mutations in the five *CSLC* glucan synthases in Arabidopsis leads to relatively mild defects (Kim *et al*., 2020). Similarly, poplar *csla* mutants lacking detectable mannans in the xylem were shown to grown normally (Guevara-Rozo *et al*., 2025). We anticipate that galactosylation and/or acetylation are critical substitutions to maintain β-mannan solubility in the Golgi and matrix polysaccharide secretion to the cell wall. The lack of enzymes or donor substrates for mannan or glucan substitution in native *Pichia* cells could explain the intracellular retention of the MannanTrack probe in glucomannan-producing yeast cells (Fig. **5a-c**, **S8**). The modularity of this platform offers further opportunities to decipher how plant GTs and GHs cooperate to define hemicellulose content and properties.

### Unraveling CSLA and MAN Roles through Yeast Synthetic Biology

While *in vitro* GH assays yield useful catalytic benchmarks and are helpful to improve enzymes for industrial use (Jacobson *et al*., 2024), they could not have predicted the unexpected biological roles of AtMAN2/5. Yeast species, including *Pichia* and *Saccharomyces* (Grieß-Osowski *et al*., 2025), can serve as surrogate hosts to characterize plant carbohydrate-active enzymes (Pauly *et al*., 2019). Intracellular MAN2/5 enzymes reduced insoluble β-mannan production by CSLAs through a variety of biochemical and microscopic assays (Fig. **4**, **5**). Surprisingly, yeast cells showed no evidence of MAN2/5 secretion even when the transmembrane domains were replaced with αMF or αGal secretion signals that worked for the MannanTrack probe and the fungal AaMAN5a enzyme (Figs. **4d**, **S3**, **S4**). Previously, MAN2 was affinity purified from *Pichia* cultures for *in vitro* assays using a similar N-terminal truncation (Wang *et al*., 2015); however, our experiments differ in the choice of the recombinant protein tags and our application of the stronger *pCAT1* promoter in episomal vectors (Vogl *et al*., 2016). Recent examples of human GTs involved in *N-*glycosylation indicate that the N-terminal transmembrane stem as well as the carbohydrate-active domains can define Golgi localization (Yagi *et al*., 2024). Amino acids beyond the predicted N-terminal membrane stem may thus contribute to the intracellular retention of MAN2/5 in yeast, although the truncation of the first 34 AtMAN2 amino acids were sufficient for its redistribution to the cytosol (Fig. **S6**), where it could no longer reduce the accumulation of insoluble mannans. The co-expression of AtMAN2/5 with CSLA elevated the content of water-soluble Man (Fig. **5d**), likely representing β-mannan oligosaccharides isolated from the endomembrane system. Using the same extraction method, *CSLC4* expression in *Pichia* produced water-soluble β-1,4-linked glucan oligosaccharides with an estimated length of four to six residues (Cocuron *et al*., 2007), while co-expression with *XXT1* (a xylosyltransferase enzyme belonging to the GT34 family that includes *MUCI10/MAGT1*) boosted insoluble glucan production (Robert *et al*., 2021).

### Potential Modes of Action for Golgi-Localized MANs

Finally, we consider three (not mutually exclusive) mechanisms by which MAN2/5 enzymes could function together with CSLAs to benefit β-mannan synthesis (Fig. **5f**): i) hydrolysis of insoluble molecules, ii) transglycosylation of oligosaccharides, and/or iii) protein interactions with CSLAs. In our yeast data, MAN2/5 enzymes partly hydrolyzed insoluble mannans made by CSLAs (Fig. **4a****, 5d**), so we favor the first hypothesis. Transglycosylation, the second mode of action, seems unlikely because purified MAN2 enzyme acted solely as a hydrolase *in vitro* (Wang *et al*., 2015). Cellulose synthases and KORRIGAN1 are known to interact (Vain *et al*., 2014), so future studies could investigate if CSLAs and MANs form protein complexes in the Golgi apparatus. Nevertheless, the catalytically inactive AtMAN2^E355A^ largely lost its hydrolytic effects despite having no obvious changes in protein expression level or subcellular localization (Figs. **S3**, **S4**), and therefore its potential for interactions with CSLA. Since crops have close orthologs of AtMAN2/5, the unexpected roles of intracellular MANs in CSLA-driven hemicellulose biosynthesis open up new paths to go glyco (Voiniciuc, 2023) and to enhance β-mannan fibers for the bioenergy and bioproducts. Even for plant MANs with characterized biochemical activities, experimental evidence for their subcellular localization is lacking (Table **S8**). Therefore, we anticipate that co-expressing CSLAs with intracellular MANs could alleviate historic bottlenecks to bioengineering plants with designer β-mannans.

## Acknowledgments

We thank the following individuals for their technical assistance: Bo Yang (initial *man2 man5* screening), Hannah Zheng (seed phenotyping), and Karen Kelley (sample preparation at the University of Florida ICBR Electron Microscopy Core Facility, RRID:SCR_019146). Sylvestre Marillonnet provided expert advice and materials for Plant MoClo assembly. Doug Smith and Mariza Miranda provided technical advice for confocal microscopy and flow cytometry at the University of Florida ICBR Cytometry Core Facility, RRID:SCR_019119. This work was supported by startup funding from UF/IFAS, the Horticultural Sciences Department, and the USDA National Institute of Food and Agriculture, Research Capacity Fund (Hatch) project 7004470. CV was also supported by core funding (Leibniz Association) from the Federal Republic of Germany and the state of Saxony-Anhalt, and by DFG grants (414353267). TJ and ME thank the PMCB program and College of Agricultural and Life Sciences (CALS) Dean’s Awards for graduate research funding. The research visit of MR was supported by a DAAD (German Academic Exchange Service) scholarship, while JM was supported by the UF University Scholars program. Finally, we thank Toshihisa Kotake for helpful discussions about MAN function since learning about our independent approaches in the summer of 2024.

## Author Contributions

TJ and CV designed the work, analyzed data and wrote the manuscript. TJ acquired most of the data, with additional contributions from ME, MQ and JM. MR and CV developed the MannanTrack yeast strain.

## Data availability

Data supporting the findings of this work are available in the main text and in the figures and tables in Supporting Information. The genetic materials generated in this current study are available from the corresponding author upon request. Popular plasmids and seeds will be donated to Addgene and ABRC, respectively. High-resolution transmission electron micrographs for yeast are on FigShare: dx.doi.org/10.6084/m9.figshare.30369646

## Supporting Information

Additional Supporting Information may be found online:

Methods S1 Supplemental Methods for Plant and Yeast Experiments

Fig. S1 Additional phenotyping of *man2 man5* seeds.

Fig. S2 Mucilage architecture after *CSLA2* overexpression in the seed coat.

Fig. S3 MAN2 and MAN5 protein structure and expression in yeast.

Fig. S4 Polymer composition following CSLA/MAN expression in *Pichia*.

Fig. S5 Fractionation of alkaline-soluble and insoluble cell wall glycans.

Fig. S6 AtMAN2 truncation alters its localization and its hydrolytic effects.

Fig. S7 Transmission electron microscopy of *Pichia* cells.

Fig. S8 Confocal microscopy of MannanTrack in yeast cells.

Fig. S9 Workflow of water-soluble mannan extraction from yeast and plant samples.

Table S1 Sequences of primers used for genotyping and quantitative PCR.

Table S2 Sequences of primers used for cloning.

Table S3 Monosaccharide composition of total mucilage extracts.

Table S4 Plant constructs and functional evaluation.

Table S5 Yeast strains used in this study.

Table S6 Summary of characterized plant MANs.

Video S1 Timelapse of Golgi-YFP and MAN2-mCherry in Tobacco

