## Supporting Information for "Golgi-Localized Mannanases Sustain Hemicellulose Biosynthesis"

Article title: Golgi-Localized Mannanases Sustain Hemicellulose Biosynthesis

The following Supporting Information is available for this article:

**Methods S1** Supplemental Methods for Plant and Yeast Experiments

**Fig. S1** Additional phenotyping of *man2 man5* seeds.

**Fig. S2** Mucilage architecture after *CSLA2* overexpression in the seed coat.

**Fig. S3** MAN2 and MAN5 protein structure and expression in yeast.

**Fig. S4** Polymer composition following CSLA/MAN expression in *Pichia*.

**Fig. S5** Fractionation of alkaline-soluble and insoluble cell wall glycans.

**Fig. S6** AtMAN2 truncation alters its localization and its hydrolytic effects.

**Fig. S7** Transmission electron microscopy of *Pichia* cells.

**Fig. S8** Confocal microscopy of MannanTrack in yeast cells.

**Fig. S9** Workflow of water-soluble mannan extraction from yeast and plant samples.

**Table S1** Sequences of primers used for genotyping and quantitative PCR.

**Table S2** Sequences of primers used for cloning.

**Table S3** Monosaccharide composition of total mucilage extracts.

**Table S4** Plant constructs and functional evaluation.

**Table S5** Yeast strains used in this study.

**Table S6** Summary of characterized plant MANs.

**Video S1** Timelapse of Golgi-YFP and MAN2-mCherry in Tobacco

**SI References**

**Methods S1 Supplemental Methods for Plant and Yeast Experiments**

**Plant Growth**

Plants were grown in walk-in growth chambers in 2.5” square deep pots or similar multiwell inserts with Jolly Gardener Pro-Line HydraFiber C/G substrate, under constant LED lights (approximately 135 µmol $m^{-2}$ $s^{-1}$), temperature (20°C) and humidity (50-60%). Dry seeds were harvested by shaking mature plants into individual paper bags.

Seed germination assays were carried out following an established protocol (Yang *et al.*, 2021), with minor adjustments. Around 30 seeds per replicate plant from multiple high-quality seed lots were placed in water and stratified in the dark at 4°C for 72 h prior to beginning the assay. Seeds were defined as germinating at the first sign of radicle protrusion.

**RNA Isolation and DNA Construct Assembly**

RNA was isolated from five mature green siliques from the main stem of 5-week-old Arabidopsis with RNeasy® Plant Mini Kit (Qiagen, MD, USA) using RLC buffer. DNase treatment was conducted on-column utilizing RQ1 DNase I (Promega, WI, USA). 200 ng RNA was used for ReverseAid First Strand cDNA Synthesis Kit (Thermo Fisher Scientific, MA, USA) with random hexamer primers. Real time PCR was quantified using 2 µL of 1:1.5 diluted cDNA containing SsoAdvanced Universal SYBR Green Supermix and analyzed on a CFX Duet Real-Time System (Bio-Rad, CA, USA).

DNA constructs were assembled using primers listed in Table S2 and the following Addgene toolkits: MoClo Toolkit #1000000044 from Sylvestre Marillonnet, MoClo Plant Parts Kit # 1000000047 from Nicola Patron), and the GoldenPiCS Kit #1000000133 from the Gasser/Mattanovich/Sauer Group. *AtMAN* coding sequences were codon optimized for *Pichia pastoris* using Benchling, synthesized by Twist Bioscience without their native stop codons to enable C-terminal fusions and initially cloned into the GoldenPiCS BB1_23 vector (Prielhofer *et al.*, 2017). Once assembled, BB1_12 vectors were used to build a single transcriptional unit into BB3rN_14 containing the *pFDH1* promoter, the *mRuby2* C-terminal tag, and *sCYCtt1* terminator elements. Final constructs were linearized with SgsI (ThermoFisher) for integration into the *Pichia* genome. AtMAN2 and AtMAN5 constructs were transformed via electroporation into WT (X-33) cells or strains harboring pPICZ B + AtCSLA2 or AkCSLA3 constructs (Voiniciuc *et al.*, 2019), or the genome-integrated BB3rN + MannanTrack probe (Robert and Voiniciuc, manuscript in preparation). Constructs were verified using colony PCR, and Sanger sequenced using vector and gene-specific primers listed in Table S4.

Subsequently, PCR was used to introduce Plant MoClo adapters and clone the MAN cDNA sequences into pAGM1287 Level 0 vectors (Engler *et al.*, 2014). The putative *pAtMAN2* and characterized *pAtTBA2* (Tsai *et al.*, 2017) promoters were PCR amplified from the Arabidopsis genomic DNA and introduced in the pICH41295 Level 0 vector. The *pUBQ10* was amplified from the pGreen20 plasmids (ABRC; CD3-2834) and domesticated by removing an internal BpiI recognition site (Pratt *et al.*). Level 1 transcriptional units were assembled into either pICH47732 or pICH47742. The FAST-RFP selection marker (Shimada *et al.*, 2010) was amplified from pICSL70008 with compatible 4-base pair recognition sites for cloning into pICH47732 and pICH47742. Level M multimeric vectors into pAGM8031 with the compatible seed coat selection marker. To avoid removing multiple BpiI and BsaI recognition sites from the genomic AtMAN2 and AtMAN5 sequences, *pUBQ10*-driven genomic sequences were PCR amplified and assembled into the final Level M constructs via HiFi cloning (NEB).

For *Pichia* episomal expression, the pPAP002 vector (Püllmann *et al.*, 2021) provided a Level 1 vector compatible with Plant MoClo full length *MAN* and *mCherry2* Level 0 parts. *AtMSR1* was amplified from an existing vector (Voiniciuc *et al.*, 2019) and cloned into Plant MoClo Level 0 acceptor (pICH41308). To inactivate AtMAN2, the "GAG" codon that encodes for E355 was changed to "GCT" via site-directed mutagenesis and MoClo assembled into Level 0 Acceptor (pAGM1287). Transmembrane regions were predicted by ARMAMEMNON (Schwacke *et al.*, 2003), compared to previous studies, and removed by amplifying the coding sequences starting with the $34^{th}$ and $33^{rd}$ codons of AtMAN2 and AtMAN5, respectively (Wang *et al.*, 2014). The αMF signal peptide was added to the sequence by generating a unique 4-base fusion site compatible with the truncated MANs unique adaptor for MoClo assembly into pAGM1287. All Level 0 parts were directly assembled into pPAP002 with either no tag, mCherry2, or FLAG C-terminal fusions.

For the *Saccharomyces* galactomannan digestion assay, an L1Sc_1F plasmid able to accept a complete Plant MoClo-compatible transcriptional unit was constructed by replacing the pAGT572_Nemo Gal-inducible pGAL1 promoter and tDIT1 terminator with BpiI-flanked lacZ cassette of the pICH47732 vector. Microbial mannanase *Aspergillus aculeatus* MAN5a cDNA (GenBank: AB015509.1) was synthesized in a *Saccharomyces*-optimized form by Twist

Bioscience with an N-terminal αGal secretion signal, C-terminal 6xHis tag with stop codon and adapters/cut sites compatible and Plant MoClo-compatible adapters. AtMAN2 and AtMAN5 cDNA with the N-terminal transmembrane portion replaced with either an N-terminal αMF or αGal- secretion signal and with the addition of a C-terminal 6xHis-tag including a stop codon were amplified from *Pichia* constructs via PCR. All constructs were cloned into L1Sc_1F with the constitutive pTDH3 promoter and the tTDH1 terminator.

### ***Pichia*** **Growth and Fluorescent Measurements**

All *Pichia* cultures were grown in 24-well plates at 30°C and 250 rpm using 1.5% methanol for induction and previously described liquid media (Robert *et al.*, 2021) with minor modifications, such as the inclusion of 150 µg/mL hygromycin (Hyg) to select for the episomal plasmid. Unless otherwise specified, three independent colonies were grown in YPD to accumulate biomass for 48 h and then transferred to YPM for 24 h to express the recombinant proteins. For Fig. S7, three independent colonies were grown for 72 h in a 24 well plate with 2 mL of yeast-peptone based media supplemented with 0.5% glycerol and 1.5% methanol (YPM+G) to stimulate biomass accumulation and recombinant protein expression.

Optical density and fluorescence were quantified using a BioTek Synergy H1 Microplate Reader (Agilent Technologies, CA, US). Fluorescence measurement centered around the following Excitation (Ex) and Emission (Em) wavelengths: CFP (Ex: 434 nm; Em: 474 nm); GFP (Ex: 480nm; Em: 518nm); YFP (Ex: 500 nm; Em: 541 nm); and RFP (Ex: 553 nm, Em: 591 nm). All measurements were taken with the gain set to 75 and used to select the three independent colonies for cell wall analyses.

For flow cytometry, cells were filtered through 35 µm strainer caps and analyzed with an Accuri C6 Flow Cytometer (BD Biosciences, USA). Measurements were taken using slow flowrate for a total of at least 10,000 cells. Data analyses were conducted in FlowJo (BD Biosciences, USA) using no fluorescence controls, RFP only and GFP only cells to select appropriate placement for the Quadrant Tool to quantify different cell subpopulations.

### **AZCL-Galactomannan Assay for MAN Activity**

For each genotype, three independent *Saccharomyces* colonies were grown in 3 mL of selection media (-uracil) with 2% Glc for 48 h at 30°C, 250 rpm in a 24-well deep plate. For each yeast strain, 750 µL of the culture mixture or of the supernatant obtained after centrifugation was aliquoted into a 1.5mL Eppendorf tube. Serial dilutions of commercial *Aspergillus niger* $\beta$-1,4-mannanase (Megazyme E-BMANN) were prepared in the yeast selection medium. To each sample, 250 µL of 0.2% insoluble AZCL-Carob Galactomannan (Megazyme I-AZGMA) in 100 mM sodium acetate was added and reactions were incubated at 37°C, 250 rpm for 48 h. The reaction tubes were centrifuged at 3,000 g for 10 min and absorbance readings of the supernatant were taken at 590 nm in a 96-well plate.

**Electron Microscopy of Seeds and Yeast**

For scanning electron microscopy (SEM), Arabidopsis seeds were transferred to conductive carbon adhesive tabs adhered to aluminum slotted 12 mm stubs, gold-palladium with argon gas sputter coated (Denton DeskV, Moorestown, NJ, USA). The specimens were examined by secondary electrons (SE), 5kV on a FE-SEM (SU-5000; Hitachi High Technologies America, Schaumburg, IL, USA).

For conventional transmission electron microscopy (TEM), *Pichia* cells after three days of culture in YPM + G medium (Robert *et al.*, 2021) were suspended in Trump's Fixative (Electron Microscopy Sciences, Hatfield, PA) for 24 h at 4°C. The fixed samples were centrifuged at 7,500 rpm for 3 min (Fisher Scientific Micro-Centrifuge Model 59A) and buffer washed with 0.1 M sodium cacodylate (pH 7.24). Cell pellets were encapsulated with low-gelling temperature agarose, type VII-A (Sigma-Aldrich, St. Louis, MO), post-fixed with buffered 2% $OsO_4$, water washed and dehydrated in a graded ethanol series (25% to 100% in 5% increments) followed by 100% anhydrous acetone. Dehydrated samples were infiltrated in graded acetone-Embed/Araldite epoxy resin with Z6040 embedding primer (Electron Microscopy Sciences, Hatfield, PA) at 30%, 50%, 70%, and 100% and cured at 70°C.

For high-pressure freezing and freeze substitution, aliquots of the same cells were pelleted at 7,500 rpm for 3 min at a time (Fisher Scientific Micro-Centrifuge Model 59A) and washed 0.1M sodium cacodylate (pH 7.24). The cells were placed into a HPM100 3 mm, 200 type-A aluminum specimen carrier (Leica Microsystems, Vienna, Austria), and prefilled with a 1-hexadecene cryoprotectant. The cells-containing specimen carriers were covered with a second type-A aluminum specimen carrier and high-pressure frozen using HPM 100 (Leica Microsystems, Vienna, Austria). The frozen samples were transferred into cryo-vials filled with anhydrous acetone containing 1% (w/v) osmium tetroxide and 0.1% uranyl acetate. Freeze substitution (FS) was carried out in a FS unit (AFS2, Leica Microsystems, Vienna, Austria) at -90°C for 72 h, -50°C for 24 h and -20°C for 24 h. During -20°C, the specimens were separated from the carriers and washed in anhydrous acetone. The FS program completed at 4°C whereas resin infiltration steps were carried out at room temperature. Embed/Araldite epoxy resin with Z6040 embedding primer (Electron Microscopy Sciences, Hatfield, PA) at 30%, 50%, 70%, and 100% infiltration steps and cured at 70°C.

For both conventionally prepared and high-pressure frozen samples, semi-thick sections (500 nm) were collected onto Fisherbrand plus slides and stained with toluidine blue. Ultra-thin 100 nm sections were collected on 200 mesh Formvar/carbon-coated copper grids and counterstained with 2% aqueous uranyl acetate and Reynold's lead citrate. Sections were examined with a FEI Tecnai G2 Spirit Twin TEM (FEI Corp., Hillsboro, OR) operated at 120 kV and digital images were acquired with a Gatan UltraScan 2k x 2k camera and Digital Micrograph software (Gatan Inc., Pleasanton, CA). All unprocessed images can be found on FigShare: dx.doi.org/10.6084/m9.figshare.30369646

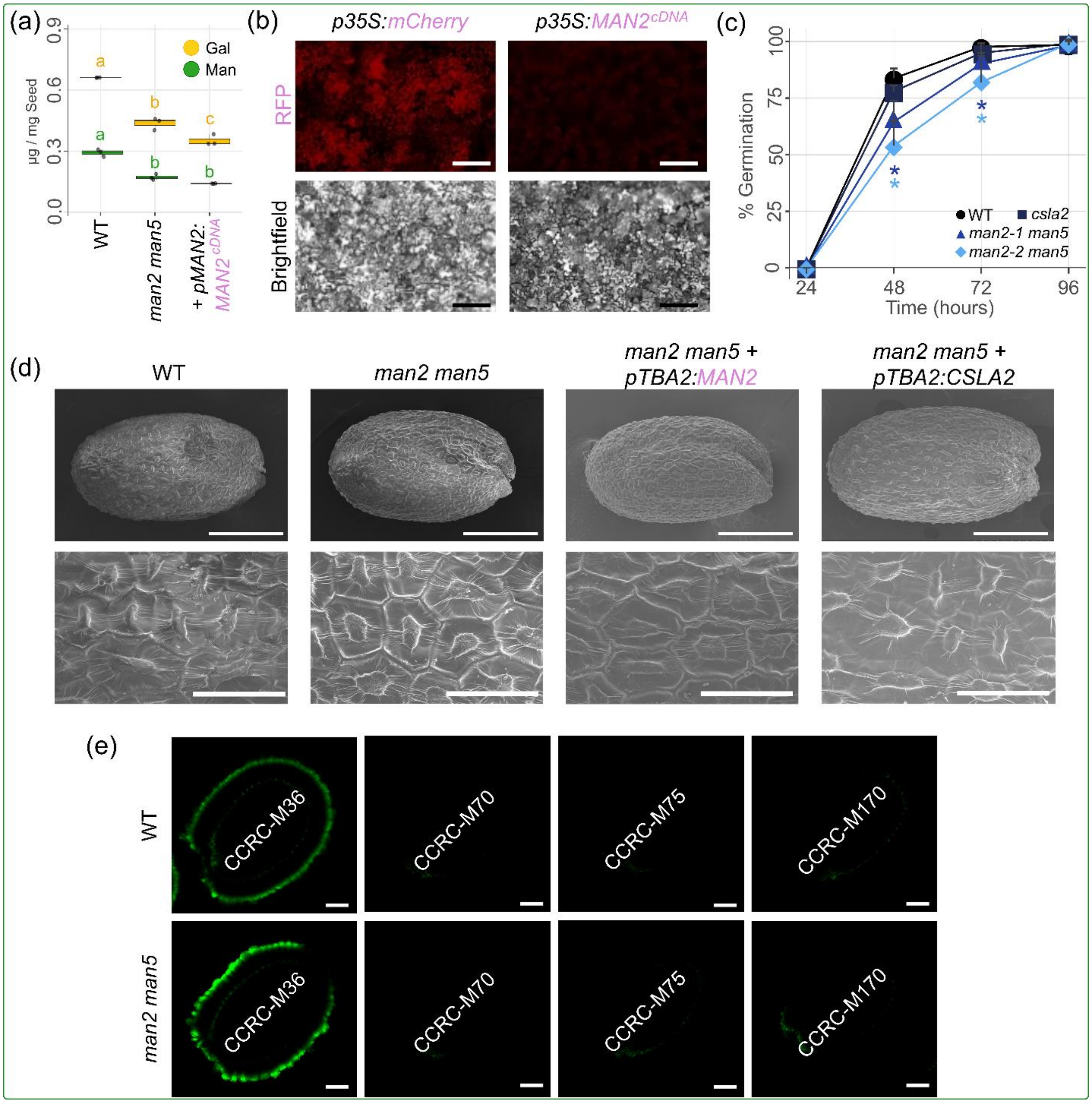

**Fig. S1** Additional phenotyping of *man2 man5* seeds. **(a)** Absolute composition of galactosylated mannans in total mucilage extracts. Data shows three biological replicates and different letters denote significant changes with one-way ANOVA (Tukey's post-hoc test, $P < 0.01$). **(b)** *N. benthamiana* leaves at 4 days post-infiltration. Red epifluorescence was captured with an Axiocam 202 mono on a Zeiss AXIO Vert A1 microscope. **(c)** Germination rates (mean + SD of three biological replicates) in water from an independent seed batch. Asterisks mark significant decreases relative to WT (Student's *t* test, $P < 0.05$). **(d)** SEM of dry seeds and seed coat epidermal cell morphology. **(e)** Seed immunolabeled with antibodies specific to pectin (CCRC-M36), and β-mannans that are galactosylated (CCRC-M70 and CCRC-M75) or acetylated (CCRC-M170). Bars: **(b)** 200 µm, **(d)** 200 µm on top row, 50 µm on bottom row, **(e)** 100 µm.

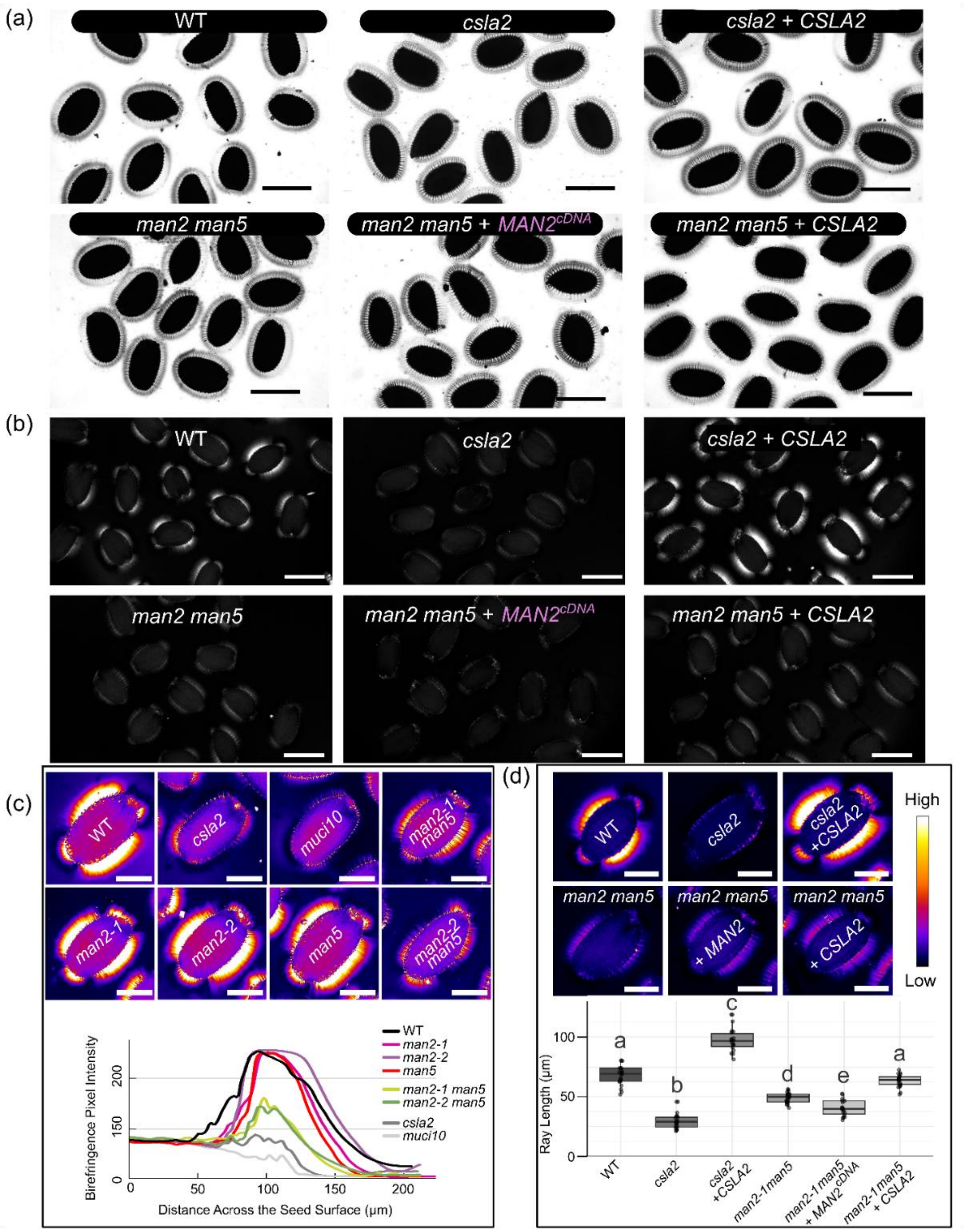

**Fig. S2** Mucilage architecture after *CSLA2* overexpression in the seed coat. Transgenes were expressed using the *pTBA2* promoter. Monochrome images of **(a)** ruthenium red staining and **(b)** mucilage birefringence. **(c)** and **(d)** Thal look up table applied to birefringence. **(c)** Relative intensity plots across the seed surface and mucilage capsules. **(d)** Birefringence ray length from 20-30 rays of respective genotypes. Different letters denote significant changes with one-way ANOVA (Tukey's post-hoc test, $P < 0.01$). For all images, magenta colored labels indicate proteins are tagged with mCherry. Bars: **(a)** and **(b)** 400 μm, **(c)** and **(d)** 200 μm.

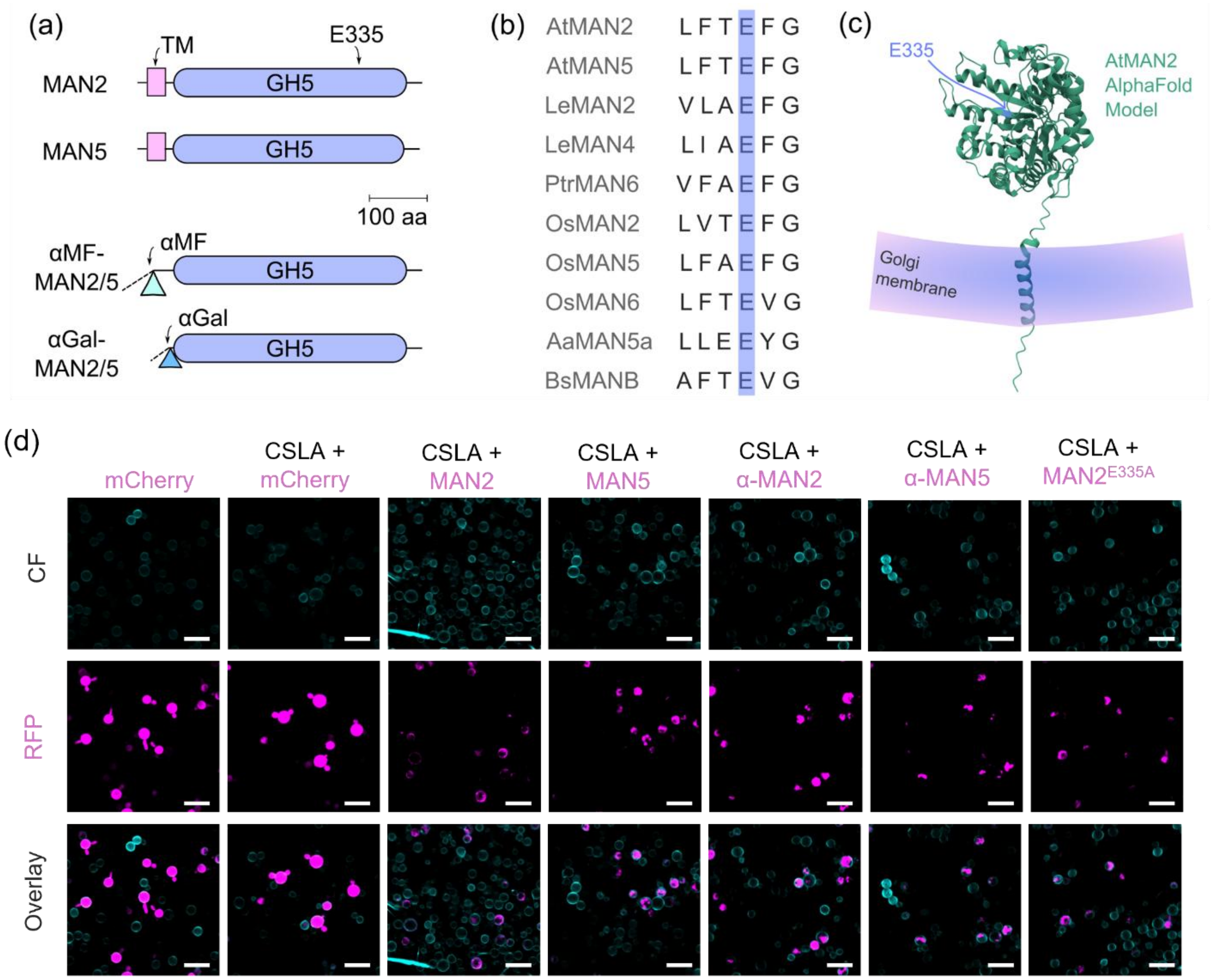

**Fig. S3** MAN2 and MAN5 protein structure and expression in yeast. **(a)** AtMAN proteins that were expressed in yeast. C-terminal tags are not shown for simplicity. **(b)** Glutamate residue at position 335 (E335) in AtMAN2 is conserved in plant and bacterial MANs. Species abbreviations: *Arabidopsis thaliana, At; Solanum lycopersicum* (formerly *Lycopersicon esculentum*), *Le; Poplar trichocarpa, Ptr; Oryza sativa, Os; Aspergillus aculeatus, Aa; Bacillus subtilus, Bs*. **(c)** AtMAN2 (Uniprot Q7Y223) folding predicted by AlphaFold. An arrow points to the position of E335 in the catalytic core, while the transmembrane domain is likely anchored in the Golgi membrane. **(d)** Optical sections of *Pichia* cells expressing mCherry-tagged MAN proteins in the untagged AkCSLA3 background. CF was used as a cell wall counterstain. Magenta labels indicate mCherry tag. Bars: **(d)** 10 µm.

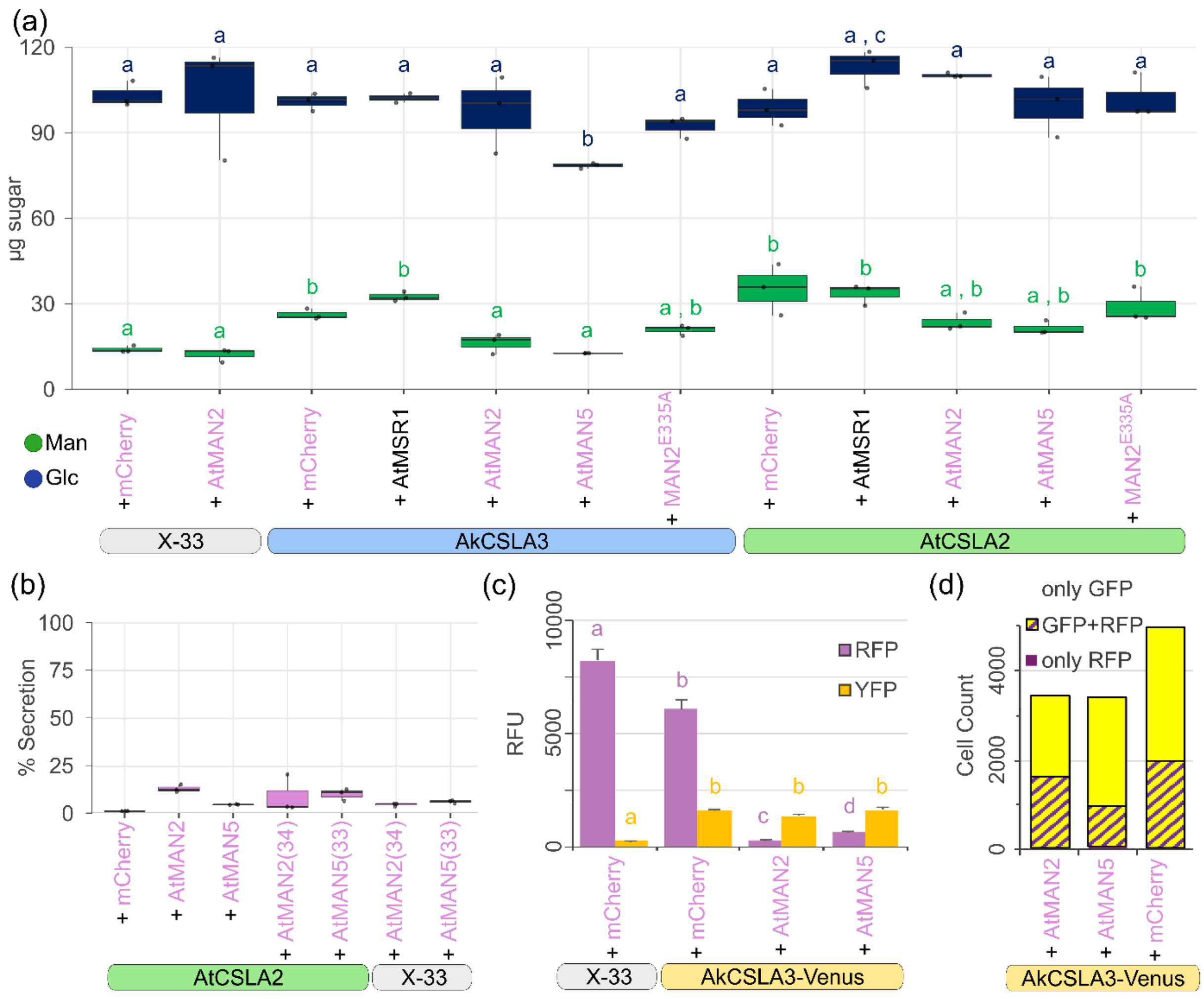

**Fig. S4** Polymer composition following CSLA/MAN expression in *Pichia*. **(a)** Absolute monosaccharide levels in equal aliquots of AKI polymers. Data show three independent colonies per genotype and correspond to the samples in Fig. 4a. **(b)** Relative secretion of mCherry fluorescence from yeast cells into the media supernatant. **(c)** Fluorescent measurements of *AkCSLA3-Venus* + *AtMAN-mCherry* yeast cultures. Data show mean + SD of three independent colonies. **(d)** Flow cytometry confirmed that only a portion of AkCSLA3-Venus cells express mCherry or tagged MANs from episomal vectors. **(a)**, **(b)**, and **(c)** different letters denote significant changes with one-way ANOVA (Tukey's post-hoc test, $P < 0.05$). For all images, magenta colored labels indicate proteins are tagged with mCherry.

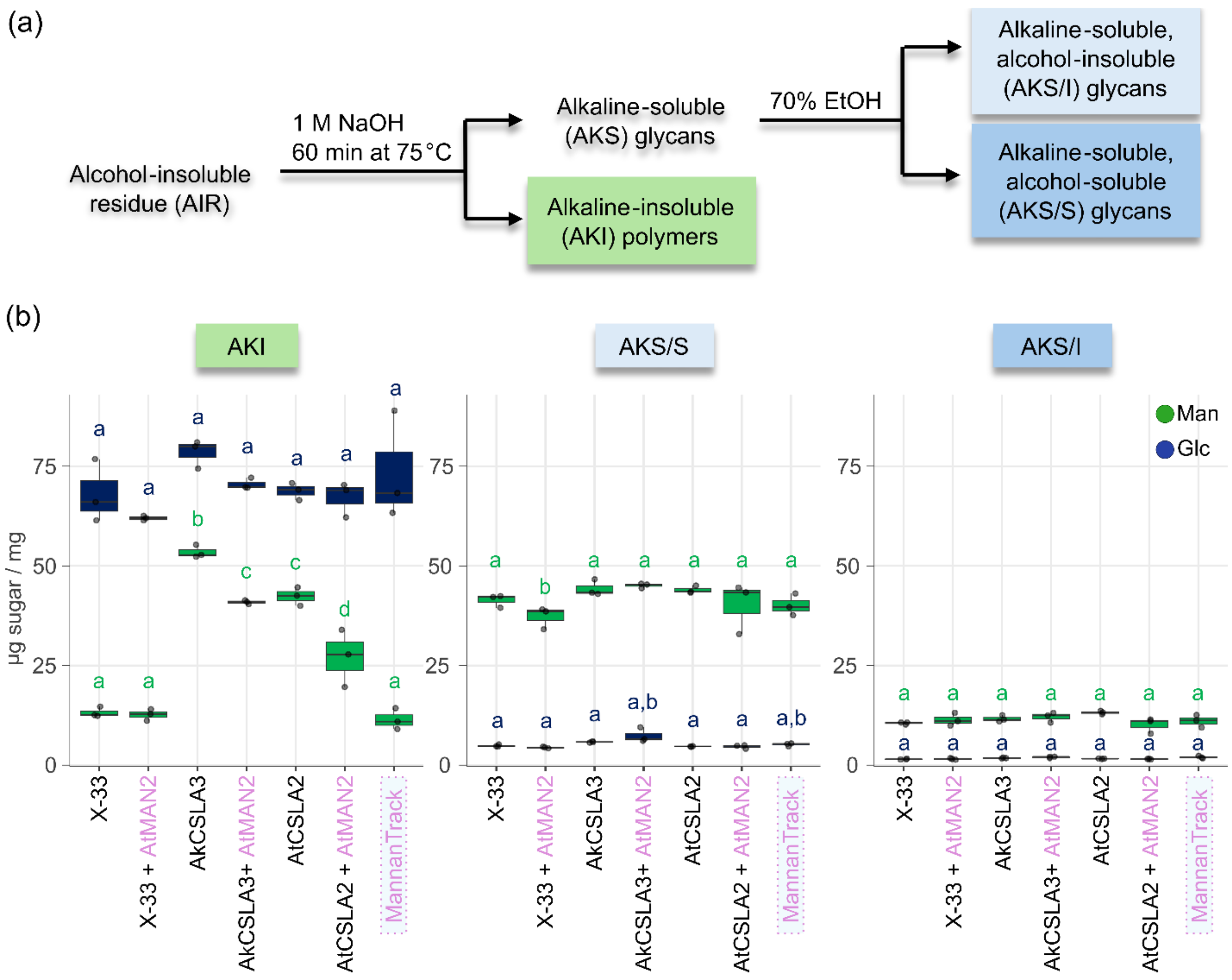

**Fig. S5** Fractionation of alkaline-soluble and insoluble cell wall glycans. **(a)** Methodology for *Pichia* cell wall fractionation adapted from (Voiniciuc *et al.*, 2019). **(b)** Composition of the three major fractions. Data show µg sugar / mg of dry weight of fraction of three biological replicates. The X-33 + MAN2 and MannanTrack strains are included as negative controls, to show that the expression of the respective proteins did not alter yeast carbohydrate metabolism. Magenta colored labels indicate proteins are tagged with mRuby2. Different letters denote significant changes with one-way ANOVA (Tukey's post-hoc test, $P < 0.01$).

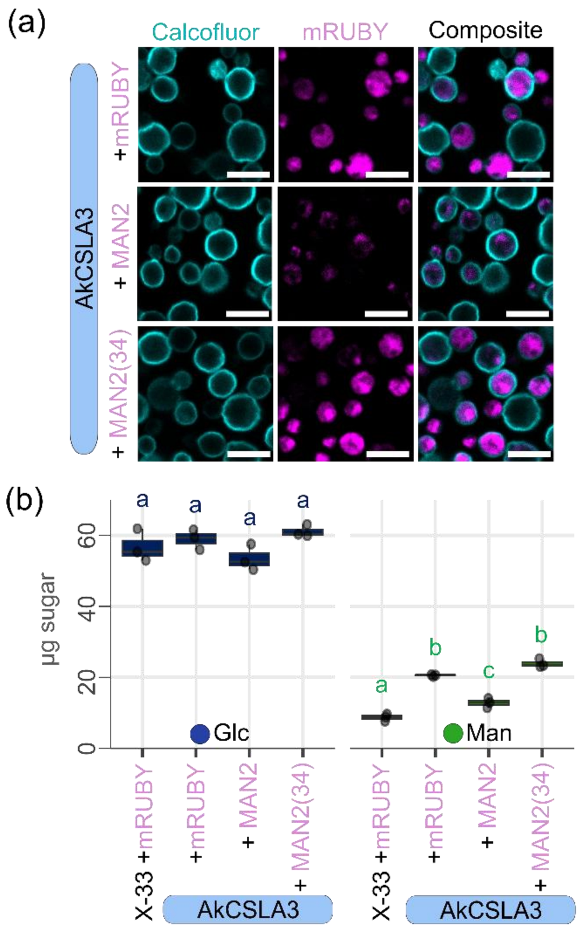

**Fig. S6** AtMAN2 truncation alters its localization and its hydrolytic effects. **(a)** MAN2(34) without its transmembrane domain or a replacement signal peptide becomes cytosolic similar to mRuby2 control expression in AkCSLA3 background. Confocal images were acquired with an 86x (1.2 NA) objective. **(b)** Biochemical effects of MAN2 truncation for three biological replicates. Different letters denote significant changes with one-way ANOVA (Tukey's post-hoc test, $P < 0.01$). Magenta labels indicate proteins are tagged with mRuby2 and expressed using the *pFDH1* promoter in an integrative vector. Bars: **(a)** 5 µm.

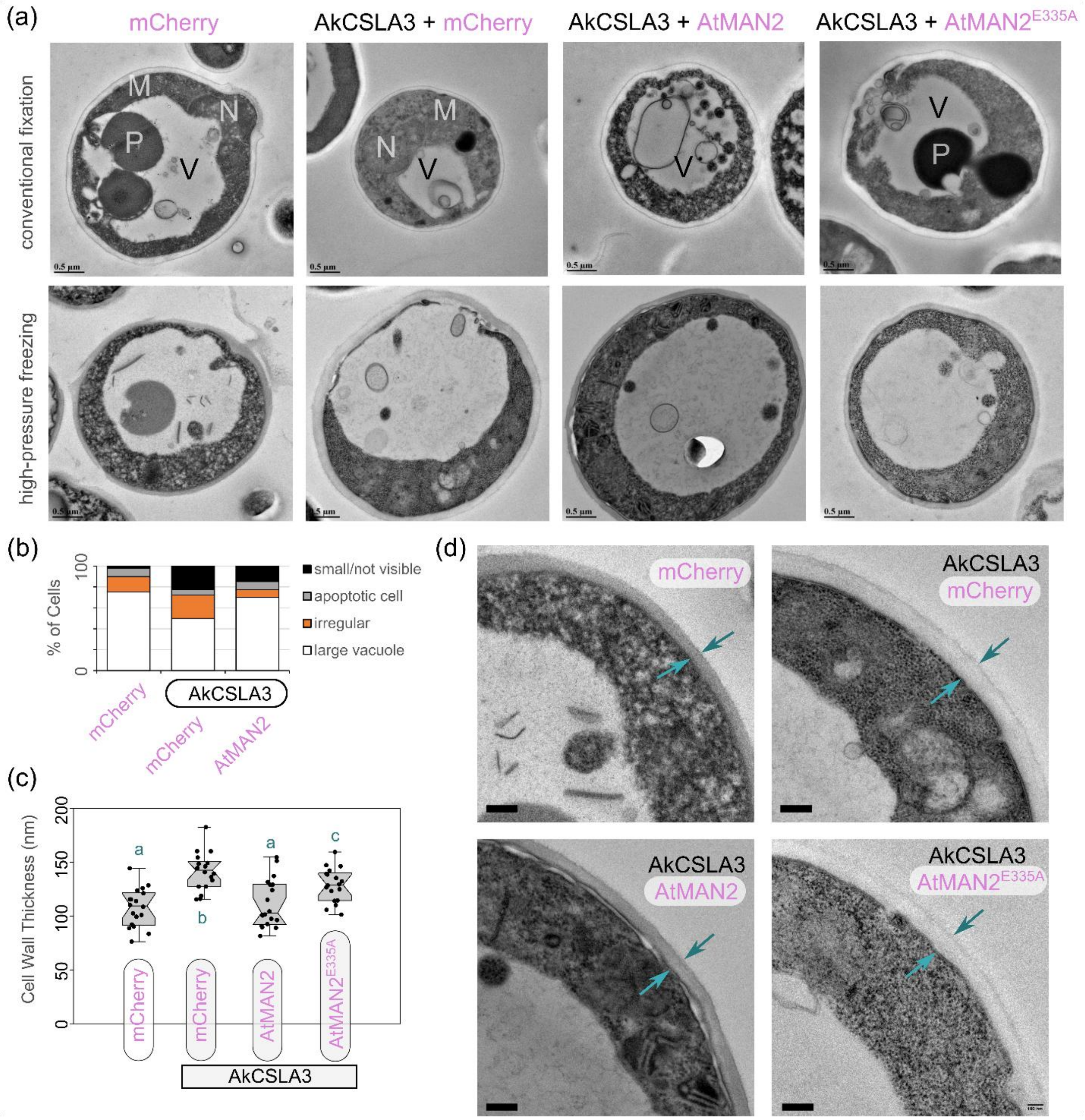

**Fig. S7** Transmission electron microscopy of *Pichia* cells. (a) Cells were imaged after conventional fixation as well as high-pressure freezing after 3 d of growth in YPM + G medium. (b) Most of the cells showed large vacuoles but no clear Golgi stacks. Vacuolar morphology (with or without autophagic bodies) was classified for 40 cells per genotype after high-pressure freezing based on a prior study using methanol induction (Vanz *et al.*, 2012). (c) Wall thickness increased in cells accumulating glucomannan and preserved with high-pressure freezing. Different letters denote significant changes with one-way ANOVA (Tukey's post-hoc test, $P <$ 0.01). (d) Arrows indicate the distance across the cell wall and magenta labels denote proteins tagged with mCherry and expressed via an episomal vector. Bars: 100 nm. High-resolution raw images from high pressure freezing and conventional fixation can be found on FigShare: dx.doi.org/10.6084/m9.figshare.30369646.

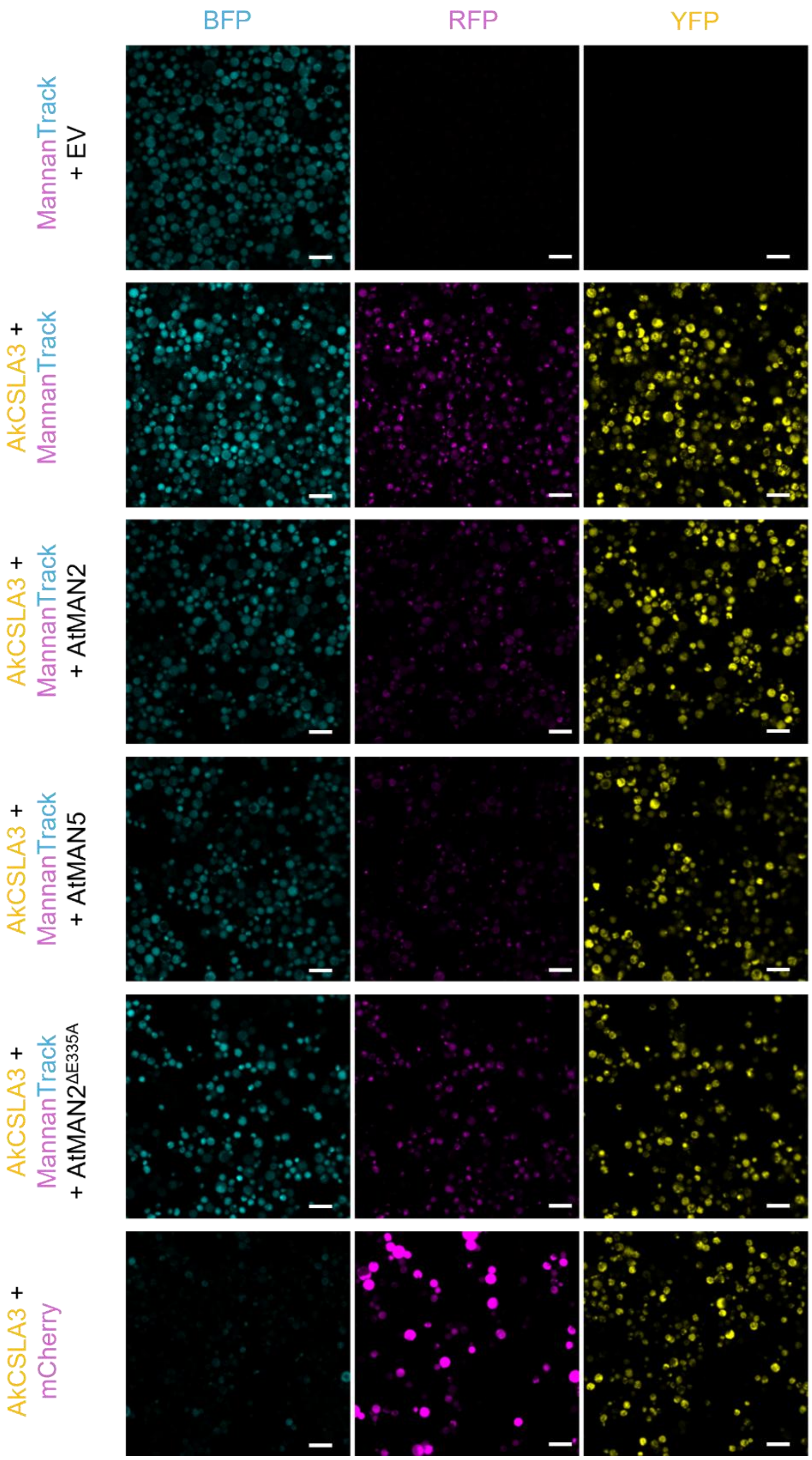

**Fig. S8** Confocal microscopy of MannanTrack in yeast cells. BFP channel corresponds to mTurquoise and serves as a cytosolic control for MannanTrack cells programmed to secrete PaCBM35a-mScarlet. RFP indicates the specific labeling of β-mannans in intracellular compartments that overlap with the location of AkCSLA3-Venus enzymes. Bars = 10 µm.

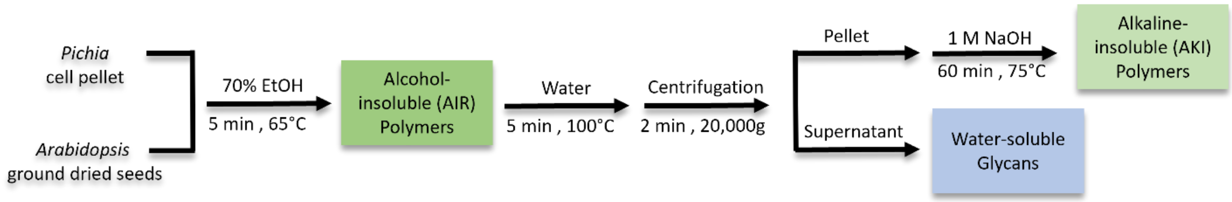

**Fig. S9** Workflow of water-soluble mannan extraction from yeast and plant samples. To quantify the insoluble mannans, we isolated AKI from *Pichia* cell pellets and AIR from *Arabidopsis* ground seeds (without mucilage extraction). This fractionation scheme corresponds to the data shown in Fig. 5d and Fig. 5e.

**Table S1** Sequences of primers used for genotyping and quantitative PCR. Primers are sorted based on their primary purpose. F and R denote the forward and reverse orientation for individual sequences or pairs.

| | Target | Forward (5' to 3') | Reverse (5' to 3') |
|---|---|---|---|
| **Plant Genotyping** | SALK LBb1.3 | attttgccgatttcggaac | |
| | GABI-KAT T-DNA | ataataacgctgcggacatctacatttt | |
| | *man2-1 (SALK_126628c.20.a)* | tatgcccatatgagaggcaag | attttcatatgccaggtgctg |
| | *man2-2 (SALK_016450)* | attttcatatgccaggtgctg | aggttgtgaacaaatgaacgg |
| | *man5-1 (707G06.01.a)* | gtgtgtgaagcagtgaagcag | tgcccactgttctggattaac |
| | *man5-2 (SALK_015220)* | gttgggctcacaattcacatc | catgaaccaatacgagttccac |
| | *man5-3 (SALK_068367)* | cgccagtgaagaaaagtatcg | agtttgtggtgaaatggatgc |
| **qPCR** | *AtMAN2* | aatcactggacgccatcttc | atgcccatatgagaggcaag |
| | *AtMAN5* | gtgaaatgggcatggcaagaagg | tcttcctggtgagcagaaccttg |
| | *AtUBQ5* | gacgcttcatctcgtcc | ccacaggttgcgttag |
| | *AtGAPC1* | tcagactcgagaaagctgctac | gatcaagtcgaccacacgg |
| | *AtCSLA2* | attccgtcggtactccaaggtc | tagcccttcctgcctcaaacag |
| | *AtMUCI10* | tacgctgcgtttcgtgaggaac | acgaacggtctccgtttgcttc |
| **Construct Genotyping** | AtMAN2 Codon Optimized | | accatgagttttggcctctg |
| | AtMAN5 Codon Optimized | | gctccagcttcaagcatttc |
| | pAtTBA2 Promoter | caacaatggtgtgttatgaaatcc | |
| | pCaMV35S Promoter | gaggagcatcgtggaaaaag | |
| | pAtMAN2 Promoter | attgtgaaaccgaacccaac | |
| | pAtUBQ10 Promoter | aacggctggatcttatgacg | |
| | mCherry | cacctacaaggccaagaagc | acatgaactgaggggacagg |
| | tAtOCS Terminator | | ttaggtttgaccggttctgc |
| | Seed coat marker | cgtgaacaaccaccacttca | |
| | Plant MoClo Level 1 | ctgatgggctgcctgtatcg | catgcacatacaaatggacgaacg |
| | Plant MoClo Level M | gctcgtcagtatgggtttgg | ggcacatacaaatggacgaac |
| | pPpCAT1 Promoter | ttgctcatcaggccaaaatc | |
| | pPpFDH1 Promoter | ttcagtgtgctgacctacacg | |
| | BB1 Integrative Pichia Vector | tgtaaaacgacggccagt | caggaaacagctatgac |
| | BB3 Integrative Pichia Vector | tcttgatccggcaaacaaac | agtcacatcatgcccctgag |
| | AkCSLA3 | tgcatctcccgacaaccatc | tcccgatctccaacaaaccg |
| | AtCSLA2 | cgccggaatatggagaatag | gttggcctgtgatctccatc |
| | AtMSR1 | tcttgtcggaaccgttgaag | tcatcaccaggtttgcttcc |
| | pScTDH3 Promoter | | tcgtaggtgtctgggtgaac |
| | tScTDH1 Terminator | | tattttaaccgccgacgttc |
| | α-mating factor | | tgccgtttcatcttctgttg |
| | α-galactosidase factor | | tttctttgaaaggtgtttttgg |
| | AaMAN5a Codon Optimized | agctatcggggtacagatgc | |
| | Saccharomyces Vector | ctgtgctccttccttcgtt | agatcacactgcctttgctg |

**Table S2** Sequences of primers used for cloning. Primers are sorted based on the technique used for DNA assembly. Golden Gate fusion sites are shown in bold.

| | Target | Forward (5' to 3') | Reverse (5' to 3') |
|---|---|---|---|
| **Golden Gate** | pTBA2 | ctggaagacta**GGAG**gttttgtgttccctacatc | gctgaagaccg**CATT**cttttgattgtttttatcttttgg |
| | pAtMAN2 pt.1 | ctcgaagactc**GGAG**atgtgggattcacatggttg | cctgaagactg**tgCC**gacagtgaaaagtaaataatacttgttg |
| | pAtMAN2 pt.2 | gacgaagacga**GGca**ctcttctcttctgtagtattagtattatgg | ctagaagacga**CATT**gctatgtggagactggtttc |
| | pUBQ10 pt.1 | gctgaagacga**GGAG**tcgacgagtcagtaataaac | gcggaagacca**gtGt**tcgatctaagattaacagaatc |
| | pUBQ10 pt.2 | gacgaagacgc**aCac**gattttctgggtttgatc | catgcgaagaccg**CATT**ctgttaatcagaaaaactcagat |
| | α-mating factor | gcggaagacgc**AATG**agaatgagatttccttcaatttttactg | gctagaagacgg**CGAC**gcttcagcctctcttttctc |
| | AtMAN2 (33) | ggcgaagactc**GCTG**aaaacggagggagag | catgaagacat**CGAA**tgtggtctgtgggaaca |
| | AtMAN5 (34) | cgagaagacgc**GCTG**aaagggaaagccaagtta | gatgaagacgt**CGAA**tgatgtctgtgagaacacag |
| | AtMSR1 | cgagaagacgg**AATG**ggtgttgatttgaggc | gtcgaagacg**AAGC**tcagcaaaagcatgaataag |
| | AtMAN2 (E335A) pt.1 | atagaagacgc**AATG**gcggctcctaca | tctgaagactag**CGGT**aaacagaacaggcttc |
| | AtMAN2 (E335A) pt.2 | gatgaagact**AAGC**ctgtggtctgtgggaacacatttca | tatgaagacta**CGAA**gttggtctgtgggaacaca |
| | MAN5 for *Sc* | ccggaagaccc**AGCC**aaagggaaagccaagttag | gatgaagacgt**CGAA**tgatgtctgtgagaacacag |
| | MAN2 for *Sc* | ccggaagaccc**AGCC**aaaacggagggagagttag | catgaagacat**CGAA**tgtggtctgtgggaaca |
| | 6xHis-tag | cggaagacgc**TTCG**catcatcatcatcatcattga | cggaagacgc**AAGC**tcaatgatgatgatgatgatg |
| | pTDH3 | cggaagacgc**GGAG**tcattatcaatactgccatttc | cggaagacgg**CATT**tttgtttgtttatgtgtg |
| | tTDH1 | ccggaagaccg**GCTT**taactcgagataaagcaatcttg | ccggaagacat**AGCG**cgttcagggtaatatattttaac |
| | α-galactosidase factor | tctgaagactc**AATG**tttgctttttatttc | ccggaagaccc**GGCT**ccaaaaacacctttc |
| **HiFi** | Backbone 1 for MAN5 | cctatggaaaaacgccagcaacgc | cactgtatctgaaactctgttaatcagaaaaactcag |
| | MAN5 gDNA | ctgattaacagagtttcagatacagtgaaactaaataac | tgctcaccatatgtctgtgagaacaaagttctttc |
| | Backbone 2 for MAN5 | tcacagacatatggtgagcaagggcg | tgctggcgtttttccataggctccgcc |
| | Backbone 1 for MAN2 | gagcctatggaaaaacgccagcaacgc | caggatacacgcacttctgttaatcagaaaaactcagat |
| | MAN2 gDNA | ctgattaacagaagtgcgtgtatcctgattcttac | ccttgctcaccattggtctatgggaacacatttcta |
| | Backbone 2 for MAN2 | ccatagaccaatggtgagcaagggcg | tgctggcgtttttccataggctccgcc |

**Table S3** Monosaccharide composition of total mucilage extracts. Data show the mean µg sugar / mg seed ± SD for three biological replicates. (a) Data corresponds to Fig. 2A. (b) Data corresponds to Fig. S2C, and cDNA sequences were driven by *pTBA2*. (c) Data corresponds to Fig. 3d. *pUBQ10* was used for the transgenes except *AtMAN5*$^{gDNA}$, which contained its native promoter.

| **(a)** | **WT** | ***csla2-3*** | ***muci10-1*** | ***man2-1*** | ***man2-2*** | ***man5-1*** | ***man2-1 man5-1*** | ***man2-2 man5-1*** |
|---|---|---|---|---|---|---|---|---|
| **Gal** | 0.48 ± 0.03 | 0.3 ± 0.01 | 0.28 ± 0.01 | 0.50 ± 0.03 | 0.51 ± 0.02 | 0.48 ± 0.05 | 0.37 ± 0.04 | 0.40 ± 0.01 |
| **Ara** | 0.19 ± 0.02 | 0.22 ± 0.00 | 0.22 ± 0.01 | 0.21 ± 0.00 | 0.21 ± 0.01 | 0.22 ± 0.03 | 0.20 ± 0.02 | 0.21 ± 0.01 |
| **Glc** | 0.35 ± 0.04 | 0.30 ± 0.10 | 0.25 ± 0.00 | 0.41 ± 0.09 | 0.41 ± 0.03 | 0.46 ± 0.15 | 0.44 ± 0.27 | 0.27 ± 0.01 |
| **Rha** | 7.42 ± 0.87 | 8.72 ± 1.25 | 9.31 ± 0.78 | 9.41 ± 2.92 | 9.41 ± 0.39 | 6.53 ± 2.91 | 7.44 ± 1.67 | 9.66 ± 0.42 |
| **Xyl** | 0.44 ± 0.11 | 0.42 ± 0.05 | 0.41 ± 0.06 | 0.38 ± 0.01 | 0.38 ± 0.02 | 0.40 ± 0.04 | 0.38 ± 0.14 | 0.47 ± 0.02 |
| **Man** | 0.27 ± 0.03 | 0.16 ± 0.01 | 0.15 ± 0.01 | 0.27 ± 0.03 | 0.27 ± 0.01 | 0.25 ± 0.04 | 0.18 ± 0.01 | 0.19 ± 0.00 |
| **GalA** | 10.42 ± 1.00 | 12.16 ± 1.85 | 12.81 ± 1.12 | 13.05 ± 3.82 | 13.05 ± 0.49 | 9.06 ± 4.01 | 12.87 ± 3.62 | 13.44 ± 0.60 |

| **(b)** | **WT** | ***man2 man5*** | ***man2 man5* + *AtCSLA2*$^{cDNA}$** | ***man2 man5* + *AtMAN2*$^{cDNA}$** | ***csla2-3*** | ***csla2-3* + *AtCSLA2*$^{cDNA}$** |
|---|---|---|---|---|---|---|
| **Gal** | 0.79 ± 0.02 | 0.52 ± 0.01 | 0.65 ± 0.09 | 0.49 ± 0.08 | 0.54 ± 0.09 | 1.06 ± 0.06 |
| **Ara** | 0.21 ± 0.00 | 0.20 ± 0.00 | 0.18 ± 0.02 | 0.20 ± 0.01 | 0.31 ± 0.06 | 0.22 ± 0.04 |
| **Glc** | 0.61 ± 0.06 | 0.51 ± 0.00 | 1.02 ± 0.14 | 0.55 ± 0.05 | 0.91 ± 0.50 | 1.06 ± 0.20 |
| **Rha** | 9.43 ± 0.80 | 8.39 ± 2.81 | 3.83 ± 0.67 | 9.48 ± 1.72 | 10.73 ± 0.82 | 8.00 ± 0.16 |
| **Xyl** | 0.99 ± 0.10 | 1.02 ± 0.08 | 0.83 ± 0.14 | 1.00 ± 0.24 | 1.07 ± 0.10 | 1.13 ± 0.04 |
| **Man** | 0.30 ± 0.05 | 0.16 ± 0.02 | 0.36 ± 0.05 | 0.14 ± 0.00 | 0.13 ± 0.02 | 0.57 ± 0.04 |
| **GalA** | 11.06 ± 0.94 | 9.91 ± 3.10 | 4.91 ± 0.87 | 11.35 ± 1.90 | 12.44 ± 1.10 | 9.85 ± 0.15 |

| **(c)** | **WT** | **WT + *mCherry*** | ***man2 man5*** | ***man2 man5* + *mCherry*** | ***man2 man5* + *AtMAN2*$^{gDNA}$** | ***man2 man5* + *AtMAN5*$^{gDNA}$** |
|---|---|---|---|---|---|---|
| **Gal** | 0.61 ± 0.02 | 0.59 ± 0.03 | 0.42 ± 0.01 | 0.48 ± 0.03 | 0.56 ± 0.02 | 0.61 ± 0.12 |
| **Ara** | 0.18 ± 0.01 | 0.18 ± 0.01 | 0.18 ± 0.01 | 0.21 ± 0.01 | 0.17 ± 0.01 | 0.21 ± 0.06 |
| **Glc** | 0.47 ± 0.06 | 0.44 ± 0.04 | 0.37 ± 0.06 | 0.41 ± 0.07 | 0.42 ± 0.04 | 0.64 ± 0.14 |
| **Rha** | 9.37 ± 0.25 | 7.65 ± 3.93 | 7.22 ± 2.75 | 9.29 ± 0.05 | 8.65 ± 0.82 | 6.55 ± 1.63 |
| **Xyl** | 0.82 ± 0.02 | 0.78 ± 0.06 | 0.81 ± 0.05 | 0.91 ± 0.03 | 0.83 ± 0.08 | 0.71 ± 0.03 |
| **Man** | 0.30 ± 0.02 | 0.27 ± 0.05 | 0.15 ± 0.01 | 0.20 ± 0.01 | 0.27 ± 0.01 | 0.25 ± 0.01 |
| **GalA** | 8.60 ± 0.42 | 7.28 ± 3.43 | 6.94 ± 2.35 | 8.93 ± 0.67 | 8.36 ± 0.67 | 6.59 ± 1.49 |

**Table S4** Plant constructs and functional evaluation. Transgenes were evaluated following stable transformation in Arabidopsis plants or transient infiltration of *Nicotiana benthamiana* leaves. Prior to the gDNA variants, only cDNA sequences were used for the plant DNA assembly. Abbreviations: FL, fluorescence, N/A, not applicable. The *pTBA2* is a seed coat-specific promoter and the putative *pMAN2* promoter was not previously tested.

| Construct | Promoter | DNA | Stable Transformation | Transient expression |
|---|---|---|---|---|
| pTBA2:AtCSLA2:tOCS | *pTBA2* | cDNA | ✓ Rescued *csla2* and altered *man2 man5* mucilage | N/A |
| pTBA2:AtMAN2-mCherry:tOCS | *pTBA2* | cDNA | ✗ No FL or rescue | N/A |
| pTBA2:AtMAN5- mCherry:tOCS | *pTBA2* | cDNA | ✗ No FL or rescue | N/A |
| pAtMAN2:AtMAN2-mCherry:tOCS | *pMAN2* | cDNA | ✗ No FL or rescue | N/A |
| p35S:mCherry:tOCS | *p35S* | cDNA | N/A | ✓ Cytosolic FL |
| p35S:AtMAN2-mCherry:tOCS | *p35S* | cDNA | N/A | ✗ No FL |
| p35S:AtMAN5-mCherry:tOCS | *p35S* | cDNA | N/A | ✗ No FL |
| pUBQ10:mCherry:tOCS | *pUBQ10* | gDNA | ✓ Cytosolic FL | ✓ Cytosolic FL |
| pUBQ10:AtMAN2$^{gDNA}$-mCherry:tOCS | *pUBQ10* | gDNA | ✓ Rescued *man2 man5* mucilage | ✓ FL punctae |
| pUBQ10:AtMAN5$^{gDNA}$-mCherry:tOCS | *pUBQ10* | gDNA | N/A | ✗ No FL |
| pAtMAN5:AtMAN5$^{gDNA}$:tOCS | *pMAN5* | gDNA | ✓ Rescued *man2 man5* mucilage | N/A |

**Table S5** Yeast strains used in this study. The list is sorted based on the final vector and shows the background used for transformation. Our laboratory generated the AtCSLA2 and AkCSLA3 strains (9), the AkCSLA3-Venus strain (16) and the MannanTrack probe (pFDH1:α-PaCBM35-pmScarlet-2A-mTurquoise:ScCYCtt). GoldenPiCS backbones BB3aZ and BB3rN (8) were used for genome integrations. BY4741 was the starting strain for *S. cerevisiae* assays with the L1Sc vector (16). The FL column outlines the fluorescent proteins expressed in each cell line. Yeast selection was based on Zeocin (Zeo), Nourseothricin (NTC), Hygromycin (Hyg), or uracil.

| | Construct | Background | FL | Selection |
|---|---|---|---|---|
| Golden PiCS | pFDH1:mRuby2:ScCYCtt | X-33 | RFP | Zeo, NTC |
| | pFDH1:AtMAN2-mRuby2:ScCYCtt | X-33 | RFP | Zeo, NTC |
| | pFDH1: AtMAN2-mRuby2:ScCYCtt | AkCSLA3 | RFP | Zeo, NTC |
| | pFDH1: AtMAN2-mRuby2:ScCYCtt | AkCSLA2 | RFP | Zeo, NTC |
| | MannanTrack | X-33 | RFP | NTC |
| | MannanTrack | AkCSLA3-Venus | CFP, YFP, RFP | Zeo, NTC |
| pPAP002 | pCAT1:mCherry2:tGAP | X-33 | RFP | Hyg |
| | pCAT1:mCherry2:tGAP | AkCSLA3 | RFP | Zeo, Hyg |
| | pCAT1:mCherry2:tGAP | AkCSLA2 | RFP | Zeo, Hyg |
| | pCAT1:mCherry2:tGAP | AkCSLA3-Venus | YFP, RFP | Zeo, Hyg |
| | pCAT1:AtMAN2-mCherry:tGAP | X-33 | RFP | Hyg |
| | pCAT1:AtMAN2-mCherry:tGAP | AkCSLA3 | RFP | Zeo, Hyg |
| | pCAT1:AtMAN2-mCherry:tGAP | AkCSLA2 | RFP | Zeo, Hyg |
| | pCAT1:AtMAN2-mCherry:tGAP | AkCSLA3-Venus | YFP, RFP | Zeo, Hyg |
| | pCAT1:AtMAN5-mCherry:tGAP | AkCSLA3 | RFP | Zeo, Hyg |
| | pCAT1:AtMAN5-mCherry:tGAP | AkCSLA2 | RFP | Zeo, Hyg |
| | pCAT1:AtMAN5-mCherry:tGAP | AkCSLA3-Venus | YFP, RFP | Zeo, Hyg |
| | pCAT1:AtMSR1:tGAP | AkCSLA3 | - | Zeo, Hyg |
| | pCAT1:AtMSR1:tGAP | AkCSLA2 | - | Zeo, Hyg |
| | pCAT1:αMF-AtMAN2-mCherry:tGAP | AkCSLA3 | RFP | Zeo, Hyg |
| | pCAT1:αMF-AtMAN2-mCherry:tGAP | AkCSLA2 | RFP | Zeo, Hyg |
| | pCAT1:αMF-AtMAN5-mCherry:tGAP | AkCSLA3 | RFP | Zeo, Hyg |
| | pCAT1:αMF-AtMAN5-mCherry:tGAP | AkCSLA2 | RFP | Zeo, Hyg |
| | pCAT1:AtMAN2($^{E335A}$)-mCherry:tGAP | AkCSLA3 | RFP | Zeo, Hyg |
| | pCAT1:AtMAN2($^{E335A}$)-mCherry:tGAP | AkCSLA2 | RFP | Zeo, Hyg |
| | pCAT1:AaMAN5a:tGAP | AkCSLA3 | - | Zeo, Hyg |
| | pCAT1:AaMAN5a:tGAP | AkCSLA2 | - | Zeo, Hyg |
| | pCAT1:AtMAN2-FLAG:tGAP | AkCSLA3-Venus + MannanTrack | CFP, YFP, RFP | Zeo, NTC, Hyg |
| | pCAT1:AtMAN2-FLAG:tGAP | AkCSLA3-Venus + MannanTrack | CFP, YFP, RFP | Zeo, NTC, Hyg |
| | pCAT1:AtMAN2(E335A)-FLAG:tGAP | AkCSLA3-Venus + MannanTrack | CFP, YFP, RFP | Zeo, NTC, Hyg |
| L1Sc | pTDH3:αGal-AaMAN5a:tTDH1 | BY4741 | - | Uracil |
| | pTDH3:αGal-AtMAN2:tTDH1 | BY4741 | - | Uracil |
| | pTDH3:αGal-AtMAN5:tTDH1 | BY4741 | - | Uracil |
| | pTDH3:αMF-AtMAN2:tTDH1 | BY4741 | - | Uracil |
| | pTDH3:αMF-AtMAN2:tTDH1 | BY4741 | - | Uracil |

**Table S6** Summary of characterized plant MANs. Abbreviations: *Solanum lycopersicum* (formerly *Lycopersicon esculentum*), *Le*; *Poplar trichocarpa*, *Ptr*; *Coffea arabica*, *Ca*; *Lactuca sativa*, *Ls*. The question mark (?) indicates a lack of experimental evidence.

| Gene | Tissues | Localization | Activity | Roles | References |
|---|---|---|---|---|---|
| *AtMAN1* | Leaves, roots, seeds | ? | Hydrolase; transglycosylase | ? | (Wang *et al.*, 2014) |
| *AtMAN2* | Stem, rosette leaves, roots, vascular tissue, seeds | Golgi apparatus (**this study**) | Hydrolase | Mannan Biosynthesis (**this study**) | (Iglesias-Fernández *et al.*, 2011; Wang *et al.*, 2015; Kikuchi *et al.*, 2025) and **this study** |
| *AtMAN3* | Flower & siliques | Cell wall | Hydrolase | Cadmium tolerance & accumulation | (Chen *et al.*, 2015) |
| *AtMAN4* | Low expression | ? | Predicted Hydrolase | ? | (Yuan *et al.*, 2006) |
| *AtMAN5* | Stem, seeds, siliques, flower, roots, leaves | Golgi apparatus (**this study**) | Hydrolase | Germination, cyanide tolerance, Mannan biosynthesis | (Iglesias-Fernández *et al.*, 2011; Yu & Xu, 2023; Kikuchi *et al.*, 2025) and **this study** |
| *AtMAN6* | Stem, root, petal, flower vascular tissue | Plasma membrane and/or Endoplasmic reticulum | Hydrolase | Suppresses SCW deposition | (Iglesias-Fernández *et al.*, 2011; Zhang *et al.*, 2023; Kikuchi *et al.*, 2025) |
| *AtMAN7* | Flower & siliques, vascular tissue | Cell wall | Hydrolase | Cadmium tolerance & accumulation | (Wu *et al.*, 2023) |
| *LeMAN1* | Seed (lateral endosperm) | ? | Hydrolase | Seed germination | (Nonogaki *et al.*, 2000) |
| *LeMAN2 (LeMAN5*)* | Seed (micropylar endosperm), (Pollen tube*) | ? | Hydrolase | Seed germination, Pollen tube development | (Nonogaki *et al.*, 2000; Filichkin *et al.*, 2004) |
| *LeMAN3* | Seed (lateral endosperm) | ? | Hydrolase | Seed germination | (Nonogaki *et al.*, 2000) |
| *LeMAN4a* | Fruit (skin & pericarp) | ? | Hydrolase; transglycosylase | Fruit ripening | (Schröder *et al.*, 2006; Wang *et al.*, 2009) |
| *PtrMAN6* | Vascular tissue | Plasma Membrane | Hydrolase | Suppresses SCW deposition | (Zhao *et al.*, 2013) |
| *CaMANA* | Seed | ? | Hydrolase | Seed dormancy release and mobilization | (Marraccini *et al.*, 2001) |
| *CaMANB* | Seed | ? | Hydrolase | Dormancy & mobilization | (Marraccini *et al.*, 2001) |
| *LsMAN1* | Seed endosperm | ? | Hydrolase | Mobilization of Carbon | (Wang *et al.*, 2004) |

**Video S1** Timelapse of Golgi-YFP and MAN2-mCherry subcellular movement. The Golgi marker is based on the cytoplasmic tail and transmembrane domain from GmMAN1 (α-MANNOSIDASE1; Glycine max). Six images acquired at 5 sec internals were exported as movie from Fiji at 1 frame per sec. (Scale bars, 50 μm).
