## Supplementary material for "Golgi-Localized Mannanases Sustain Hemicellulose Biosynthesis": Video S1

### Slide 1
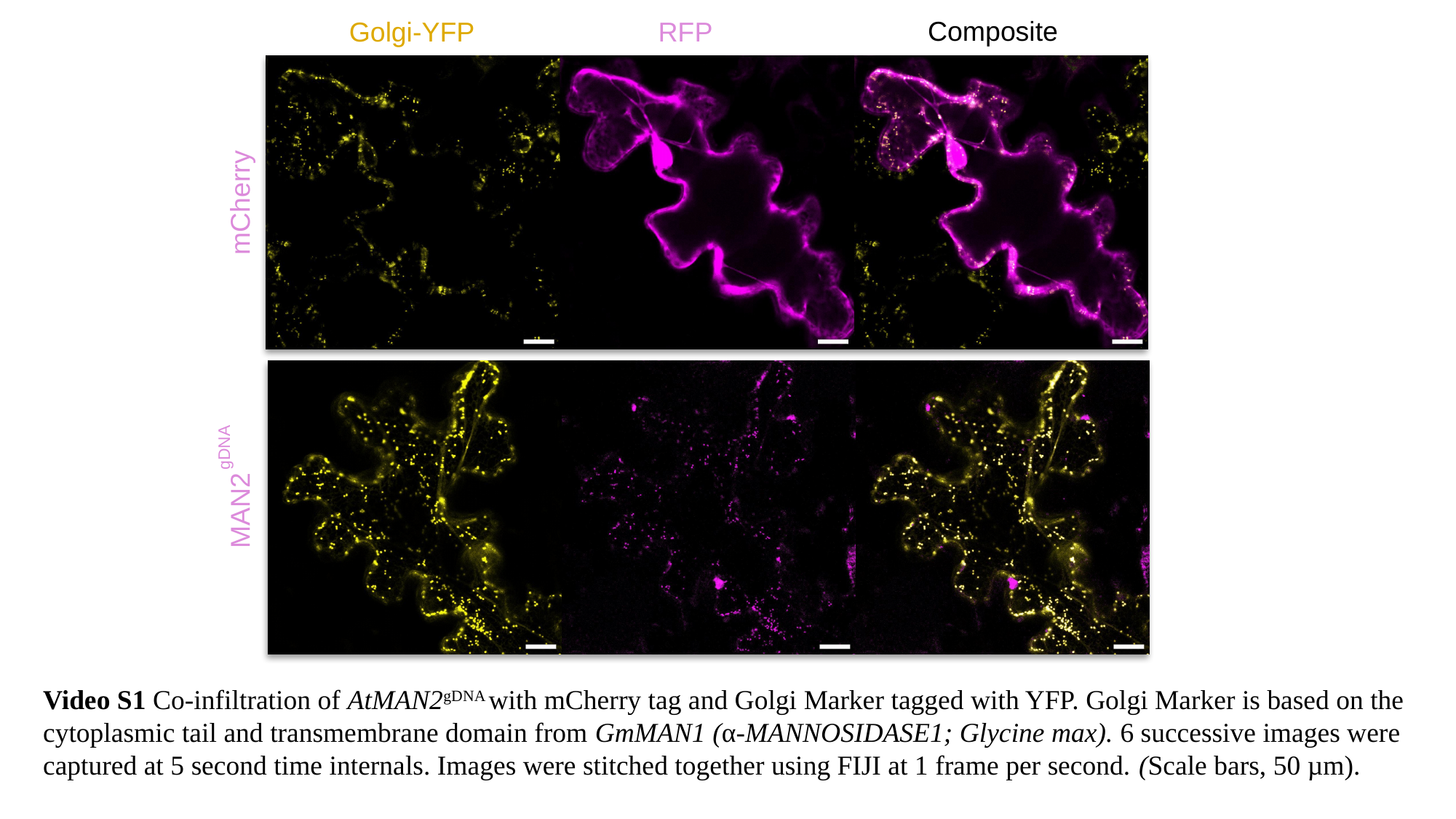

RFP
Golgi-YFP
Composite
mCherry
gDNA
MAN2
Video S1 Co-infiltration of AtMAN2^gDNA^ with mCherry tag and Golgi Marker tagged with YFP. Golgi Marker is based on the cytoplasmic tail and transmembrane domain from GmMAN1 (α-MANNOSIDASE1; Glycine max). 6 successive images were captured at 5 second time internals. Images were stitched together using FIJI at 1 frame per second. (Scale bars, 50 µm).
